# *Borrelia burgdorferi* loses essential genetic elements and cell proliferative potential during stationary phase in culture but not in the tick vector

**DOI:** 10.1101/2024.10.28.620338

**Authors:** Jessica Zhang, Constantin N. Takacs, Joshua W. McCausland, Elizabeth A. Mueller, Jeline Buron, Yashna Thappeta, Jenny Wachter, Patricia A. Rosa, Christine Jacobs-Wagner

## Abstract

The Lyme disease agent *Borrelia burgdorferi* is a polyploid bacterium with a segmented genome in which both the chromosome and over 20 distinct plasmids are present in multiple copies per cell. This pathogen can survive at least nine months in its tick vector in an apparent dormant state between blood meals, without losing cell proliferative capability when re-exposed to nutrients. Cultivated *B. burgdorferi* cells grown to stationary phase or resuspended in nutrient-limited media are often used to study the effects of nutrient deprivation. However, a thorough assessment of the spirochete’s ability to recover from nutrient depletion has been lacking. Our study shows that starved *B. burgdorferi* cultures rapidly lose cell proliferative. Loss of genetic elements essential for cell proliferation contributes to the observed proliferative defect in stationary phase. The gradual decline in copies of genetic elements is not perfectly synchronized between chromosomes and plasmids, generating cells that harbor one or more copies of the essential chromosome but lack all copies of one or more non-essential plasmids. This phenomenon likely contributes to the well-documented issue of plasmid loss during in vitro cultivation of *B. burgdorferi*. In contrast, *B. burgdorferi* cells from ticks starved for 14 months showed no evidence of reduced cell proliferative ability or plasmid loss. Beyond their practical implications for studying *B. burgdorferi*, these findings suggest that the midgut of the tick vector offers a unique environment that supports the maintenance of *B. burgdorferi*’s segmented genome and cell proliferative potential during periods of tick fasting.

**Importance:** *Borrelia burgdorferi* causes Lyme disease, a prevalent tick-borne illness. *B. burgdorferi* must survive long periods (months to a year) of apparent dormancy in the midgut of the tick vector between blood meals. Resilience to starvation is a common trait among bacteria. However, this study reveals that in laboratory cultures, *B. burgdorferi* poorly endures starvation and rapidly loses viability. This decline is linked to a gradual loss of genetic elements required for cell proliferation. These results suggest that the persistence of *B. burgdorferi* in nature is likely shaped more by unique environmental conditions in the midgut of the tick vector than by a general innate ability of this bacterium to endure nutrient deprivation.

## Introduction

The spirochete *Borrelia burgdorferi* and closely related genospecies are the causative agents of Lyme disease, the most prevalent vector-borne human disease in North America and Europe (1–4). To persist in nature, this pathogen cycles between *Ixodes* ticks and susceptible vertebrates hosts (5–10). Humans are dead-end hosts that can become infected when incidentally bitten by a *B. burgdorferi*-colonized tick (8, 11, 12). During the enzootic cycle, tick larvae acquire *B. burgdorferi* by feeding on an infected animal (5, 10, 11, 13–15). The spirochetes reside in the tick midgut during the larval molting and remain there until the nymph feeds on a host. In response to tick feeding, *B. burgdorferi* multiplies in the tick midgut before migrating to the salivary glands where it is transmitted to a vertebrate host (16–18). For *Ixodes* ticks, intervals between blood meals can often last several months to a year (19, 20) during which *B. burgdorferi* persists in the largely nutrient-devoid lumen of the tick midgut (10, 11, 14, 15, 21–24). The number of *B. burgdorferi* cells per tick remains relatively stable for at least up to 9 months (16, 19, 25), consistent with a starvation-induced growth arrest. This has led to the hypothesis that *B. burgdorferi* can withstand long periods of starvation without losing cell proliferative capability when re-exposed to nutrients; this is further supported by observations of rapid spirochete replication upon tick feeding (16, 17, 26, 27).

Many bacteria exhibit remarkable resilience to nutrient deprivation (28, 29), a prevalent condition in nature due to fierce competition and limited resources (30, 31). In fact, nongrowing bacteria are thought to be the norm rather than the exception. In the laboratory, batch stationary phase cultures are often used as models to study bacterial dormancy. Various non-sporulating bacterial species retain cell proliferative potential (i.e., ability to form colonies on agar plates containing fresh medium) for months or even years without external addition of nutrients (28, 32–36). These studies have identified that adaptation to stationary phase is commonly associated with transcriptional reprogramming, changes in cellular morphology, and genetic selection (28, 37).

*B. burgdorferi* is distinct from these commonly studied bacteria as it is an obligate parasite that cannot survive outside of a tick or vertebrate host. This bacterium has limited metabolic capacities and is auxotrophic for all amino acids, nucleotides, and fatty acids (11, 38). Despite this, *B. burgdorferi* can be cultured in a highly complex, nutrient-rich medium (BSK-II) to saturating densities around 10^8^ cells/mL at stationary phase (39–41). The defined mammalian tissue culture medium RPMI 1640, which lacks essential nutrients found in BSK-II medium, is commonly used to induce starvation of *B. burgdorferi* cultures (18, 42–47). Both in vitro stationary phase- and RPMI 1640-starved populations have been leveraged to identify *B. burgdorferi* genes involved in survival in the tick vector (10, 18, 42–44, 46, 47) and to evaluate antibiotic susceptibilities (48–51). However, we recently showed that *B. burgdorferi*, which is polyploid in exponential phase cultures, reduces the number of copies of its chromosome and essential plasmid (cp26) per cell upon entry to stationary phase (52). This raises the question of whether *B. burgdorferi* retains cell proliferative potential under starvation conditions. We aimed to address this question given that recovery from dormancy is critical for *B. burgdorferi*’s transmission and pathogenesis. Therefore, we performed a quantitative characterization of stationary phase *B. burgdorferi* cultures with respect to cell replicative potential and genome composition and compared it to the natural situation.

## RESULTS

### *B. burgdorferi* cells in stationary phase cultures lose their ability to proliferate in vitro

To characterize stationary phase phenotypes, we monitored cultures of the non-clonal *B. burgdorferi* strain B31-MI grown in BSK-II medium under common laboratory conditions (34°C in the presence of 5% CO_2_). For these experiments, we used two biological replicates (rep) that were followed for over 20 days total, with over 19 of those days occurring during stationary phase. For both cultures, we examined the cell count by dark field microscopy. Both replicates of B31-MI reached stationary phase at densities at or above 10^8^ cells/mL (Fig. 1A, black and grey circles, corresponding to rep1 and rep2). These cell densities were maintained for the remaining time of the experiment (Fig. 1A), suggesting that little to no cell lysis occurred during this period of starvation. In our hands, stationary phase cells mostly retained their characteristic flat-wave shape throughout the course of the experiment (Fig. 1B).

**Figure 1.**
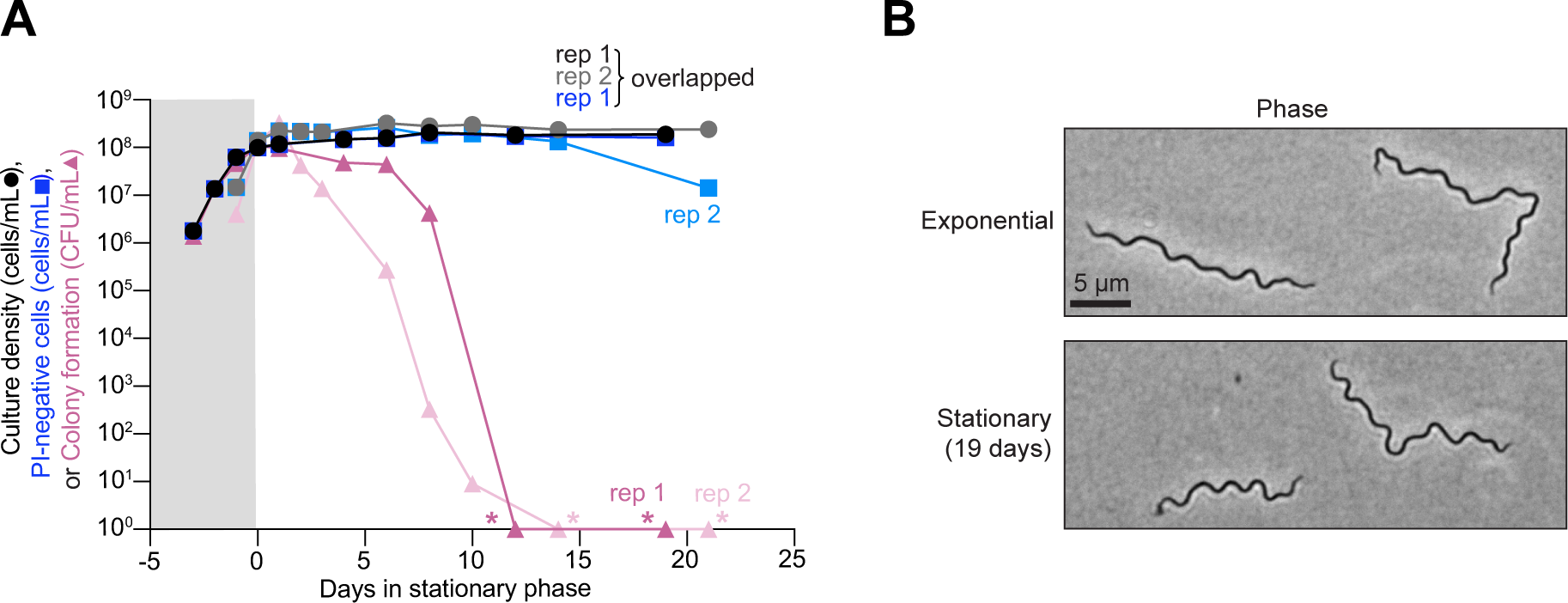
In vitro *B. burgdorferi* cultures lose cell proliferative capability in stationary phase. **A.** Results from two independent cultures (biological replicates, rep 1 and 2) of strain B31-MI. The plot shows a comparison between the cell culture density (black and grey circles, corresponding to rep1 and rep2) determined by visual direct counting, the calculated number of cells stained negative for propidium iodide (PI) (dark and light blue squares, corresponding to rep1 and rep2) as determined by fluorescence microscopy, and the ability of cells to form colonies (dark and light pink triangles, corresponding to rep1 and rep2) assessed by semisolid BSK-agarose plating using samples of the same culture. For PI-negative cell determination, 72 to 417 cells were analyzed for each strain and time point (see Supplemental File 1 for specific n values). When no colonies were detected (pink asterisks), this data was plotted as one detected colony to show the drop in CFU/mL in the log scale. Gray and white backgrounds indicate exponential and stationary phases, respectively. **B.** Representative phase-contrast images of B31-MI cells in exponential phase and after 19 days of stationary phase.

Quantitation of membrane integrity by propidium iodide (PI) staining of samples from the same cultures revealed that either 5 or 86% of cells retained intact membranes (i.e., remained PI-negative) after about 21 days in stationary phase depending on the replicates (Fig. S1A). The reason for this variability between biological replicates is unclear, but even in the lowest case, we calculated that over 10^6^ cells/mL (10^8^ cells multiplied by the PI-negative percentage at each time point) remained PI-negative in the culture after ≥ 19 days in stationary phase (Fig. 1A, dark and light blue squares) in line with a previous study (53). Thus, our calculations suggest that a large number of cells in both stationary phase cultures retained an intact membrane. In comparison, the ability of these cells to form colonies decreased far more rapidly during stationary phase (Fig. 1A, dark and light pink triangles). By ten days in stationary phase, the colony forming units (CFU) dropped by 7 to 8 orders of magnitude, after which we could no longer detect colonies (asterisks, Fig. 1A). By contrast, exponentially growing cultures readily formed colonies (Fig. 1A) (54, 55), with calculated plating efficiencies near 100% (Fig. S1B). This near-perfect plating efficiency indicates that virtually all cells in the exponential phase cultures could form colonies. The large discrepancy between CFUs and calculated PI-negative cell abundance (Fig. 1A) indicates that membrane integrity is a poor indicator of the ability of stationary phase *B. burgdorferi* cells to proliferate once plated.

We observed similar trends in reduced cell proliferative potential for three other B31 clonal derivatives (S9, CJW_Bb378, and CJW_Bb379) monitored up to seven days in stationary phase (Fig. S1C-E). Furthermore, we showed that the decrease in proliferative potential of stationary phase cells was consistent regardless of using a limiting dilution assay in liquid culture or a colony plating method (Fig. S1E, orange hexagons vs. pink triangles). Similar trends for culture density, cell proliferation, and PI-staining were observed for two other commonly studied non-clonal *B. burgdorferi* strains, 297 and N40, when monitored for up to 20 days in stationary phase (Fig. S1F-G). Altogether, our results show that *B. burgdorferi* cells in stationary phase cultures, including those with normal flat-wave morphology and intact membranes, lose their ability to proliferate upon nutrient repletion.

### Both environmental acidification and acute starvation in cultures disproportionality impact *B. burgdorferi*’s proliferative capacity relative to its membrane integrity

In stationary phase cultures, *B. burgdorferi* experiences not only starvation, but also gradual acidification of its environment, a consequence of sugar fermentation to lactic acid (38, 39, 56–59). Medium acidification occurred with all strains tested but varied in magnitude (Figs. 2A and S2A, dark and light purple diamonds). Loss of survival has previously been reported after 24 h incubation of *B. burgdorferi* cells in acidified medium (60). To expand on this study, we resuspended cells from exponentially growing cultures (density ∼10^7^ cells/mL) of strain K2 (a clonal B31 derivative) in fresh BSK-II medium acidified to pH 6.0 by addition of lactic acid prior to culture inoculation. We chose pH 6.0 as it was the lowest pH reached by stationary phase cultures across all strains tested (Figs. 2A and S2A), as shown before (53). *B. burgdorferi* cells inoculated in this acidified but nutrient-rich medium maintained relatively constant cell densities (Fig. 2B, black and grey circles), consistent with prior reports that pH 6.0 medium inhibits growth (61). Medium acidification had little effect on membrane integrity, as the calculated number of PI-negative cells remained comparatively unchanged for both replicates (Figs. 2B and S2B, dark and light blue triangles). In contrast, prolonged incubation in acidic BSK-II was associated with a gradual decrease in the ability of cells to form colonies, with the CFUs dropping by more than seven logs after 14 days of incubation at pH 6.0 (Fig. 2B, dark and light pink triangles).

**Figure 2.**
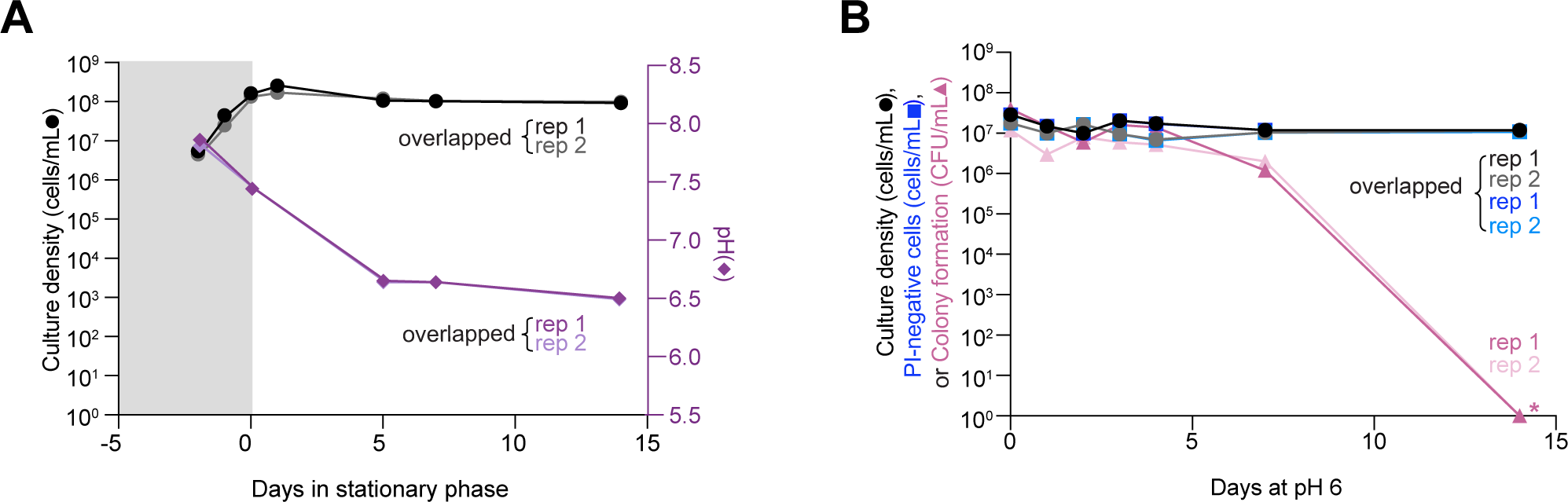
Cultures at pH 6.0 lose cell proliferative ability. Results from two independent cultures (biological replicates, rep 1 and 2) of strain K2 are shown. **A.** Plot showing culture density (black and grey circles, corresponding to rep1 and rep2) vs. pH (dark and light purple diamonds, corresponding to rep1 and rep2) for strain K2 (a clonal derivative of B31-MI) in BSK-II medium. Gray and white backgrounds indicate exponential and stationary phases, respectively. **B.** Plots showing the effects of medium acidification to pH 6.0 on the ability of *B. burgdorferi* cells to grow, form colonies after plating and maintain the integrity of their cell membrane based on uptake of propidium iodide (PI). Cells from two independent exponentially growing K2 cultures (density of ∼10^7^ cells/mL) were pelleted and resuspended into the same volume of fresh BSK-II medium pre-adjusted to pH 6.0. Culture densities expressed as cells/mL are shown as black and grey circles corresponding to replicates (rep) 1 and 2, respectively. Calculated PI-negative cells/mL values are shown with dark and light blue squares corresponding to rep 1 and 2. For PI-negative cell determination, 31 to 105 cells were analyzed for each strain and time point. CFU/mL are shown as dark and light pink triangles corresponding to rep 1 and 2, respectively. When no colonies were detected (pink asterisks), this data was plotted as one detected colony to show the drop in CFU/mL in the log scale.

Under natural conditions in the tick or mammalian host, *B. burgdorferi* is unlikely to encounter a pH low enough to become detrimental for cell proliferation (61). Therefore, we next examined the effects of prolonged starvation alone, i.e., without significant environmental acidification. To do this, we resuspended *B. burgdorferi* K2 cells from cultures with densities from 2-4x10^7^ cells/mL (i.e., before reaching stationary phase) in RPMI 1640 and incubated for several days to induce starvation in *B. burgdorferi* (18, 42, 44, 45). Consistent with prior studies (42, 62), *B. burgdorferi* cells in RPMI 1640 arrested growth with the pH of the medium remaining ≥ 7, even after 14 days of incubation (Fig. 3A). Acute starvation in RPMI 1640 resulted in the formation of membrane bulges and condensed ball-like cell forms known as round bodies (Fig. 3B, white arrowheads), as reported previously (18, 42, 44, 45, 63), The percentage of round bodies in the cell population rapidly increased, reaching a plateau of ∼90% after seven days of incubation in RPMI 1640 (Fig. 3C). These round bodies often stained PI-positive (Fig. 3B), indicating a loss of membrane integrity. Despite the increase in PI-positive cells (when considering both spirochetes and round bodies) and the corresponding decrease in PI-negative cells in the population (Fig. S3A), we again observed a large discrepancy between the fraction of PI-negative cells and the CFUs. By day 14, the CFU values of RPMI 1640 cultures had dropped by over three orders of magnitudes, compared to a single order of magnitude reduction of PI-negative cells (Fig. 3D, dark and light pink triangles). The loss of cell proliferative ability in RPMI 1640 was even more severe for strains N40 and a 297 derivative (Bb914) while the percentage of round body morphotypes and PI-negative cells in the population was similar across strains (Fig. S3A-C). Collectively, our results suggest that both starvation and pH 6.0 cause considerable loss in *B. burgdorferi’s* ability to proliferate under in vitro culture conditions.

**Figure 3.**
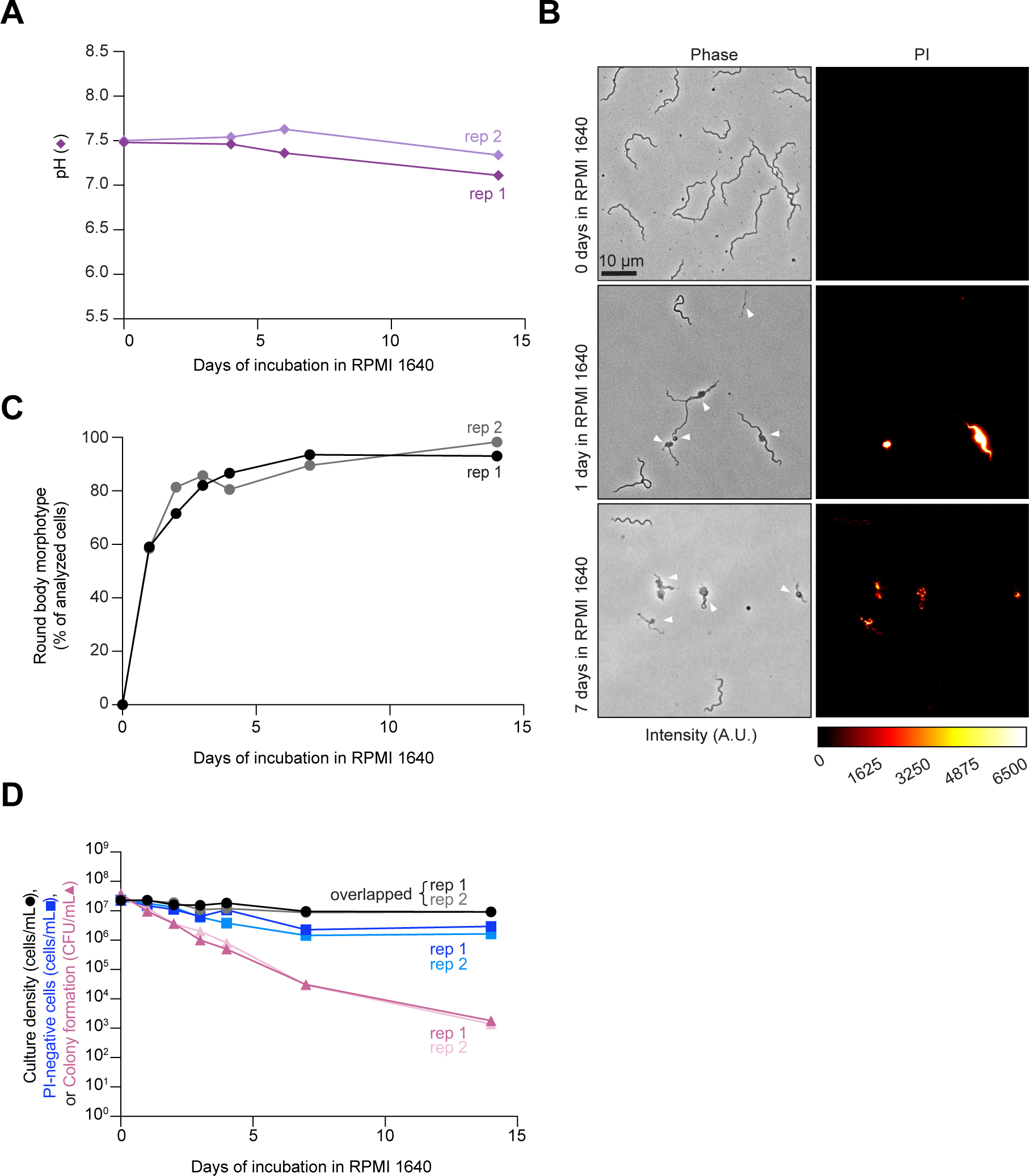
Cultures in nutrient-deficient RPMI 1640 medium lose cell proliferative ability over time. **A.** Plot showing the pH evolution of two independent cultures (biological replicates, rep 1 and 2) in which cells from exponential phase cultures of strain K2 were resuspended in RPMI 1640 medium. Dark and light purple diamonds correspond to rep 1 and 2, respectively. **B.** Representative phase contrast and fluorescent images of cells of strain K2 after the indicated days of incubation in RPMI 1640 medium. Cells were incubated with propidium iodide (PI) prior to imaging. White arrowheads point to cells with morphological defects that include membrane bulges, condensed ball-like forms, and ghost-like cells with decreased phase contrast. The color scale bar for pixel intensity values applies to all PI images shown, where A.U. indicates arbitrary units. **C.** Plot showing the percentage (%) of cells that displayed the round-body morphotype in the same cultures as in (B). For round-body determinations, 48 to 247 cells were analyzed for each strain and time point (see Supplemental File 1 for specific n values). Black and grey circles correspond to rep 1 and 2, respectively. **D.** Plot showing a comparison between the cell culture densities determined by visual direct counting (black and grey circles, corresponding to rep1 and rep2), the calculated number of PI-negative cells assayed by fluorescence microscopy (dark and light blue squares, corresponding to rep1 and rep2), and CFU/mL (dark and light pink triangles, corresponding to rep1 and rep2) assessed by semisolid BSK-agarose plating using samples of the same cultures as in (B) and (C). For PI-negative cell determination, 48 to 247 cells were analyzed for each strain and time point (see Supplemental File 1 for specific n values).

### Stationary phase is associated with loss of chromosomal material

We wondered whether the loss of cell proliferative potential in stationary phase cultures was associated with DNA changes. This question was partly triggered by the observation that the DNA staining with Hoechst 33342 shifted from a homogeneous DNA pattern throughout the cytoplasm in exponentially growing cells (52, 64) to a patchy distribution in stationary phase cells, with noticeable gaps of DNA-free areas (Fig. S4A, blue arrowheads). Quantification revealed that the percentage of B31-MI, 297, and N40 cells with homogeneous DNA staining (i.e., a single detected Hoechst-stained object) rapidly decreased in the first five days of stationary phase (Fig. S4B-D). After five to ten days in stationary phase (depending on the strain), the intracellular DNA signal became too weak (Fig. S4E-G) to reliably quantify the number of Hoechst-stained objects. This may be due to a reduction in DNA concentration and/or a decrease in cell membrane permeability to the DNA dye. Regardless, the measured mean Hoechst signal decreased exponentially with a half-life of 1-4 days depending on the strain (Fig. S4E-G).

Using a ParB/*parS* labeling system, we previously showed that the copy numbers of chromosomal origin (*oriC)* and plasmid cp26 per cell decrease when cultures transition to stationary phase (52). This loss of *oriC* copy numbers per cell was confirmed by quantitative polymerase chain reaction (qPCR) (52), validating the ParB/*parS* labeling system (52). We reproduced the stationary phase-induced decrease in *oriC* copy number per cell with two B31-derived strains in which *oriC* is labeled with chromosomally expressed mCherry-ParB (strain CJW_Bb379) (Figs. 4A and S5A). This was confirmed by labeling the *oriC* region with another chromosomally expressed protein fusion, ParZ-GFP (strain CJW_Bb378) (Figs. 4A and S5A). ParZ-GFP binds a DNA region close to *oriC* similar to mCherry-ParB (52). We also showed that the density of *oriC* copies per 10 μm of cell length decreased in stationary phase (Fig. S5B), confirming that the reduction in *oriC* density is not due to a change in average cell length in the population. This decline led to a mixed population of stationary phase cells lacking fluorescently labeled *oriC* foci (cells without yellow arrowheads, Fig. 4A) and cells with at least one labeled *oriC* (cells with yellow arrowheads, Fig. 4A). The percentage of cells lacking a clear fluorescent *oriC* focus, and thus a full chromosome, drastically increased with time in stationary phase (Fig. 4B). This likely contributes to the observed loss of cell proliferative potential, as cells would be unable to self-replicate without a copy of their essential chromosome carrying an origin of replication.

**Figure 4.**
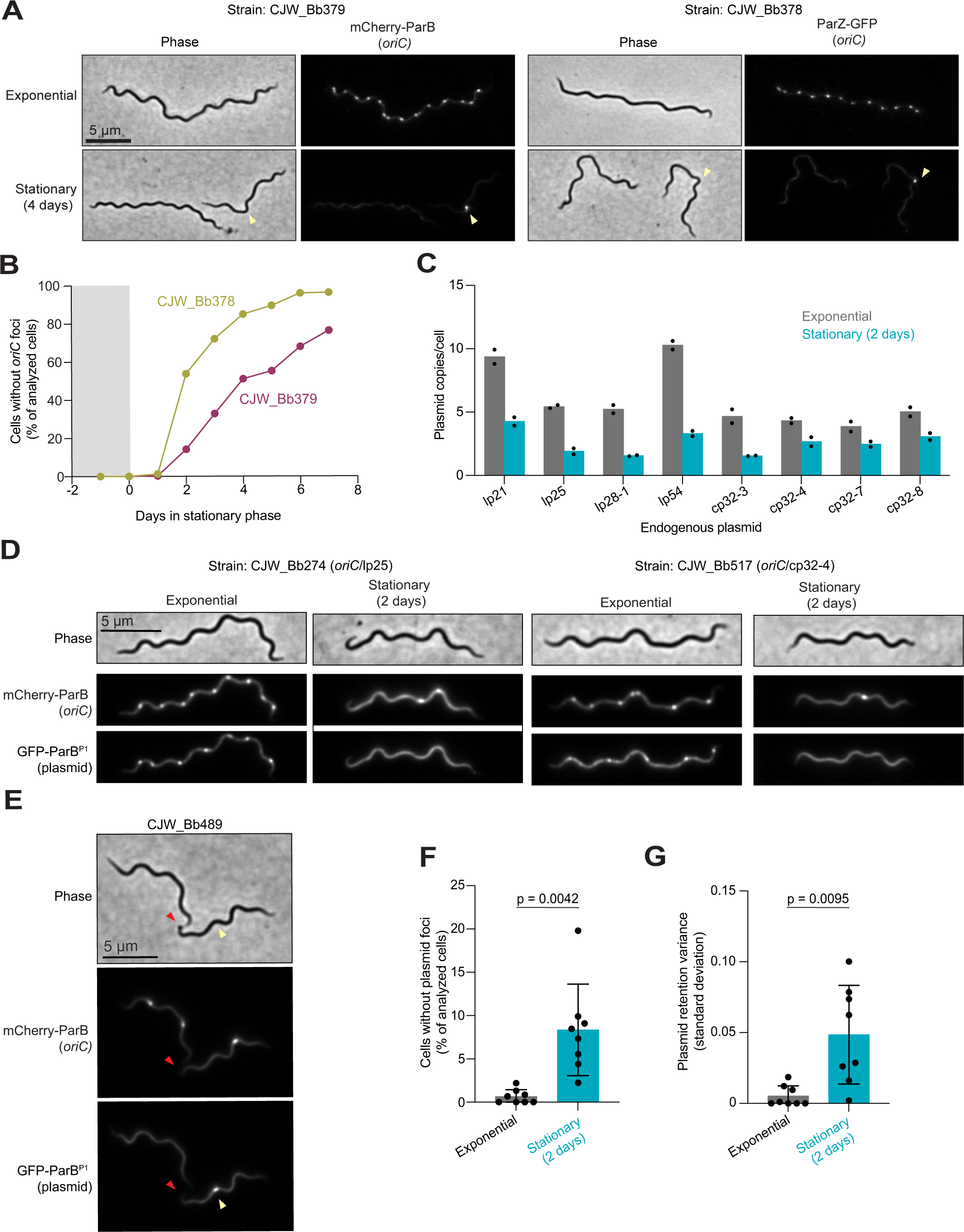
Stationary phase is associated with the generation of cells without copies of *oriC* or specific plasmids. **A.** Representative phase contrast and fluorescence images of cells of strains CJW_Bb379 and CJW_Bb378 in which the *oriC* region is labeled with mCherrry-ParB or ParZ-GFP, respectively. Images are shown for each strain during exponential phase or four days after the cultures entered stationary phase. Yellow arrowheads point to the stationary phase cell with a clear *oriC* focus. **B.** Plot showing the percentage of the cell populations without clear fluorescent *oriC* foci, using the same cultures as in (A). Gray and white backgrounds indicate exponential and stationary phases, respectively. For each strain and time point, 60 to 387 cells were analyzed (see Supplemental File 1 for specific n values). **C.** Plot showing the mean plasmid copies per cell in exponential phase (grey bars) and after two days in stationary phase (teal bars). Each black dot represents an independent biological replicate. These measurements were obtained through image analysis of cultures of eight strains expressing msfGFP-ParB^P1^ and carrying its target *parS^P1^* sequence inserted in the indicated plasmid while mCherry-ParB labeled the chromosomal *oriC* region. Only cells with at least one clear *oriC* focus were considered in this analysis. The strain identities and the number of cells analyzed for each data point (n = 22-167) are detailed in Supplemental File 1. **D.** Illustrative images of selected cells of strains CJW_Bb274 (with fluorescent markers of *oriC* and lp25) and CJW_Bb517 (with fluorescent markers of *oriC* and cp32-4) in exponential phase and after two days in stationary phase. mCherry-ParB and msfGFP-ParB^P1^ selectively label *oriC* and plasmid copies, respectively. Plasmid loss is highlighted by the lack of fluorescent foci in the selected cells. **E.** Illustrative images of a selected cell of strain CJW_Bb489 in stationary phase for two days. This cell, which carries fluorescent markers of *oriC* and lp28-1, was in the process of completing cell division to yield a daughter cell lacking a fluorescent lp28-1 focus. Yellow arrowheads point to the future daughter cell that will inherit a copy of lp28-1. The division site is indicated by red arrowheads and corresponds to decreased cellular fluorescence in the mCherry and GFP channels (see signal intensity profiles in Fig. S5D). **F.** Plot showing the percentage of the cell populations without plasmid foci in strains in which both *oriC* and a given endogenous plasmid are labeled. Results for cells in exponential phase (grey bars) are compared to those for cells after two days in stationary phase (teal bars) using the same cultures as in (C). Each black dot represents the mean of two independent biological replicates per strain (n = 22-167 cells per replicate). Bars represent the average percentage of the cell population without a specific plasmid across the eight evaluated strains, while error bars indicate standard deviations. **G.** Same as in (F) but comparing the variability (standard deviation) in plasmid retention in cells. Each black dot represents the mean of two independent biological replicates per strain (n = 22-167 cells per replicate). Bars represent the average plasmid retention variance across the eight evaluated strains, while error bars indicate standard deviations.

### Loss of plasmids in cells is associated with cultivation in stationary phase

Next, we wondered whether the well-known loss of plasmids in cell cultures (65–70) is primarily associated with stationary phase. We reasoned that plasmids that are not essential for growth in culture may also decrease their copy number during stationary phase. If so, some cells in stationary phase may lose all copies of an endogenous plasmid before losing all copies of the chromosome. If these cells experience nutrient repletion, they may be able to proliferate due to the non-essential nature of the lost plasmid in culture, resulting in a population that lacks this plasmid. To test this hypothesis, we compared the copy number of eight non-essential plasmids in exponential phase and after two days in stationary phase, using four linear plasmids (lp21, lp25, lp28-1, and lp54) and four circular plasmids (cp32-3, cp32-4, cp32-7, and cp32-8) as representatives. Therefore, we imaged strains in which both *oriC* and one of the representative plasmids are fluorescently labeled (52). We quantified the number of fluorescently labeled plasmid copies in cells that contain at least one *oriC*, excluding cells with no labeled *oriC* from the analysis to eliminate cells unable to proliferate. We found that all eight examined endogenous plasmids were present in a lower copy number per cell (or per 10 μm of cell length) after two days in stationary phase when compared to exponential phase (Figs. 4C and S5C). We also observed stationary phase cells with at least one copy of *oriC* but no apparent plasmid foci, as illustrated in Fig. 4D for plasmids lp25 and cp32-4. The generation of such cells was observed in some dividing cells in which each future daughter cell carried at least one copy of *oriC* but one of them lacked any copy of the labeled plasmid. This is illustrated for lp28-1 in Fig. 4E where the red arrowheads show the site of reduced fluorescence intensities corresponding to the division site (see intensity profiles in Fig. S5D) and yellow arrowheads point to the cell half that will inherit *oriC* but not lp28-1 after division. Quantification confirmed that cells have a greater probability of losing a non-essential plasmid in stationary phase relative to exponential phase (Fig. 4F, p = 0.0042). The variance in plasmid retention also increased in stationary phase (Fig. 4G, p = 0.0095), suggesting a dysregulation of plasmid maintenance.

We validated our hypothesis that plasmid loss increases in frequency in stationary phase by plating stationary phase cells and analyzing the plasmid complement of the resulting colonies (clones) after growth using a multiplex PCR assay (71). We determined that ∼17% (4/24) of clones isolated from 10-day-old stationary phase cultures had lost at least one endogenous plasmid (Table 1 and Fig. S6A). In contrast, all clones (22/22) isolated from the exponential growth phase of the same culture had a full plasmid complement (Table 1 and Fig. S6B). Thus, stationary phase leads to a higher likelihood of plasmid loss from cells, likely because the gradual loss of plasmid copies is not perfectly synchronized with that of chromosome copies, occasionally creating proliferation-competent cells with a reduced genome size due to plasmid loss.

**Table 1.** Plasmid retention in different growth stages measured by plasmid profiling of isolated clones of strain CJW_Bb523 (for in vitro studies) and CJW_Bb474 (for in vivo studies).

| Growth condition | Replicate | Plasmid loss summary |  | Plasmid loss details |  |
| --- | --- | --- | --- | --- | --- |
|  |  | Clones analyzed: number | Clones that lost plasmids: number (% of total) | Clone ID | Plasmids lost |
| Exponential phase culture | Culture A | 11 | 0 (0%) |  |  |
|  | Culture B | 11 | 0 (0%) |  |  |
| Stationary phase culture, day 10 | Culture A | 12 | 1 (8.3%) | A-S10-2 | lp36 |
|  | Culture B | 12 | 3 (25%) | B-S10-2 | cp32-4 |
|  |  |  |  | B-S10-3 | lp38 |
|  |  |  |  | B-S10-8 | lp56 |
| Ticks colonized for one year | Nymph 1 | 9 | 0 (0%) |  |  |
|  | Nymph 2 | 10 | 0 (0%) |  |  |
|  | Nymph 3 | 10 | 0 (0%) |  |  |
|  | Nymph 4 | 8 | 0 (0%) |  |  |
|  | Nymph 5 | 10 | 0 (0%) |  |  |

### Loss of *B. burgdorferi* plasmids is not detected in ticks unfed for over a year

*B. burgdorferi* is believed to experience growth arrest in its tick vector after blood meal digestion and molting. This conclusion is based on immunostaining and cell plating, which show that the total spirochete number, though highly variable from tick to tick, remains, on average, relatively constant in unfed, post-molt nymphal or adult ticks over 3-9 months (16, 19). This in vivo growth arrest is thought to be due to nutrient limitation (10, 15, 20, 72, 73).

In a previous study, we reported the colonization of ticks with a *B. burgdorferi* strain (CJW_Bb474) by larval stage feeding on infected 4-8 weeks-old female mice (*Mus musculus*) (52). These larvae were allowed to molt into nymphs, which were then maintained at room temperature. After a month, we determined the number of viable spirochetes in a subset (six nymphs) of the entire cohort by plating for CFUs (52). For this current study, we performed the same analysis on five nymphs from the same cohort that had remained unfed at room temperature for 14 months (Fig. 5A). While there was some variability across individual ticks (especially for the one-month time point), the spirochete loads did not decrease between one-month and 14-month unfed nymphs (Fig. 5B). If anything, the spirochete loads may slightly increase, though the statistical significance was marginal (p = 0.036). This is consistent with the presumption that spirochete growth is limited in unfed ticks.

**Figure 5.**
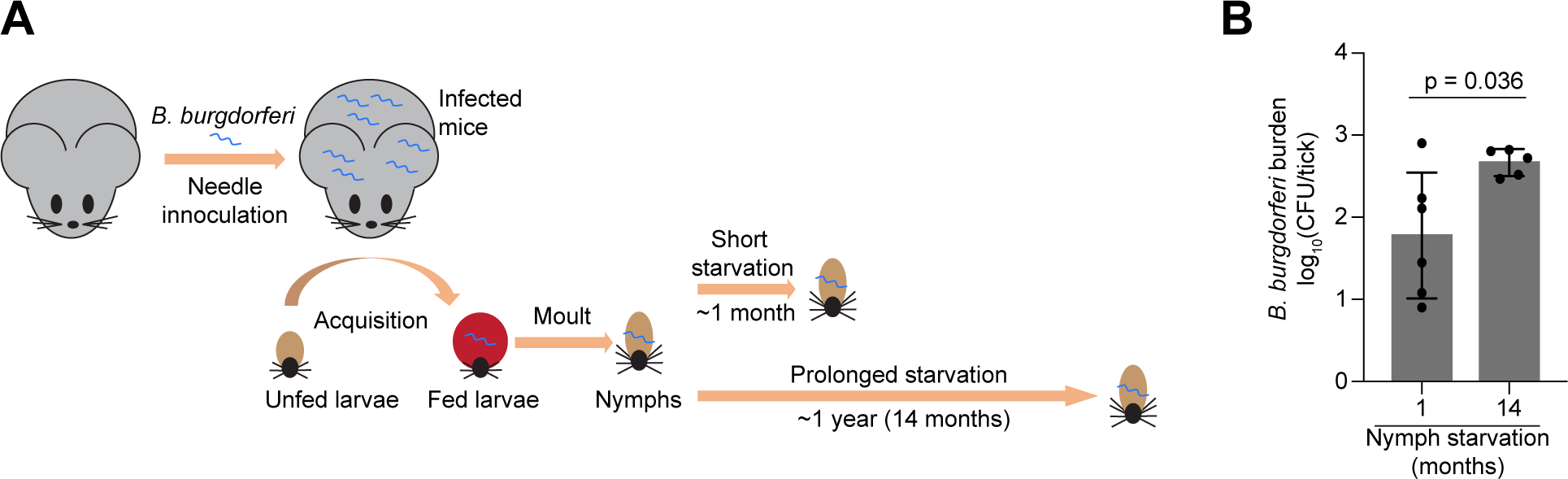
*B. burgdorferi* cells in *I. scapularis* nymphs starved for 14 months remain proliferative. **A.** Schematic of the experiment. Four to eight weeks-old female mice were infected with *B. burgdorferi* strain CJW_Bb474 by needle inoculation. Larval *I. scapularis* ticks became colonized with this strain during feeding on the infected mice. The fed larvae were allowed to molt into nymphs. These nymphs were kept unfed at room temperature for the indicated durations before their spirochete loads were determined by plating. The mouse and tick cartoons are adapted from a previous publication (52). **B.** *B. burgdorferi* burdens in unfed nymphs that were generated as described in (A). The *B. burgdorferi* burdens were measured one month or 14 months after molt by semisolid BSK-agarose plating and are expressed as log_10_(CFU/tick). Shown are means ± standard deviations for five or six individual ticks. The individual values are indicated by black dots. The unpaired *t* test with Welch’s correction on the log_10_-transformed data gave a p value of 0.036. The one-month and 14-month nymphs came from the same cohort The one-month data, which were previously published (52), are shown here for comparison.

Next, we determined the plasmid complement for 47 *B. burgdorferi* clones derived from plating the extract of five 14-month-old nymphs (8-10 clones per tick). We found that all 47 clones retained the full plasmid complement of the parental CJW_Bb474 strain (Table 1 and Fig. S7). Thus, the plasmid loss observed within days in laboratory stationary phase cultures does not occur when *B. burgdorferi* is maintained in the starved tick vector, even after a 14-month period of apparent quiescence.

## DISCUSSION

Our study shows that *B. burgdorferi* in BSK-II cultures poorly tolerates stationary phase under standard culturing conditions, as shown by the precipitous loss in CFUs within days of growth arrest (Figs. 1A and S1C-F). This contrasts with *B. burgdorferi* cells harvested from ticks starved for over a year (Fig. 5) despite using comparable plating techniques. Inside the tick, *B. burgdorferi* experiences ambient temperatures, which are often lower than our laboratory culture conditions (34°C). We found that the decrease in CFUs and *oriC* copies per cell in stationary phase was slower when *B. burgdorferi* cultures (CJW_Bb379) grew in BSK-II at ∼21°C (Fig. S8) relative to 34°C (Figs. 1A and S5A). This is not surprising given that enzymatic activities and thus metabolism are slower at lower temperatures. Importantly, the significant reduction in CFUs at ∼21°C occurred within a few weeks (Fig. S8) whereas no reduction in CFUs was observed after over a year inside tick midguts (Fig. 5B), indicating that temperature alone cannot explain the maintenance of cell proliferative capacities in unfed ticks.

We also found that stationary phase cells lost proliferative potential even when their cytoplasmic membrane appeared uncompromised based on the absence of PI staining (Figs. 1A and S1F). It has long been debated whether bacteria with apparent intact membrane integrity can remain viable even if they have lost their ability to be cultured in a medium that normally supports their growth (74, 75); this notion has been proposed for *B. burgdorferi* as well (51). Such cells are often referred to as “viable-but-nonculturable”. While the term “viable” can have different definitions, we are primarily interested in the proliferative potential of *B. burgdorferi* when nutrients become available, as *B. burgdorferi’s* enzootic cycle and pathogenesis require cell multiplication. Could stationary phase *B. burgdorferi* cells that were unable to form colonies in BSK plates be “resuscitated” under other conditions that are more physiological than the laboratory growth medium? While this is formally possible, we find this scenario unlikely for two reasons. First, *B. burgdorferi* cells that remained in starved ticks for over a year were able to self-replicate on same BSK plates, indicating that BSK plates are suitable growth media for regrowth even after long periods of growth limitation. Second, we observed loss of essential genetic elements in stationary phase cultures. This genetic loss generates cells that lack chromosomal elements required for cellular replication, such as *oriC* (Fig. 4A-B). Furthermore, we also observed stationary phase cells that have lost all copies of cp26 (e.g., cell without an arrowhead in Fig. S5E), an endogenous plasmid that carries genes (e.g., telomere resolvase-encoding gene *resT*) essential for cell proliferation (76). The loss of these essential genetic elements is accompanied by a large drop in cell proliferative potential (Fig. 1). In contrast, the average number of spirochetes in ticks did not decrease after 14 months without a blood meal (Fig. 5), which could have been expected if *B. burgdorferi* cells had lost essential genetic elements. We also found no evidence of plasmid loss in the spirochetes recovered from these ticks (Fig. S7 and Table 1).

The rapid loss of proliferative potential in *B. burgdorferi* stationary phase cultures contrasts with what is observed with a variety of other bacteria, including but not limited to *Escherichia coli*, *Mycobacterium smegmatis, Sarcina lutea*, and *Serratia marcescens* (28, 29, 31–36, 77, 78). This is likely due to the limited metabolic activity of *B. burgdorferi* in comparison to these free-living bacteria. Collectively, our data indicate that the tick midgut provides a favorable environment for *B. burgdorferi*’s survival over long periods of apparent growth arrest. While medium acidification to pH 6.0 affects *B. burgdorferi* proliferative capability in vitro (Fig. 2B), such low pH is unlikely to be encountered in the tick host. The midgut of unfed ticks maintains a pH of ∼7.4 (similar to the pH 7.6 of fresh BSK II medium) even when colonized with *B. burgdorferi* (61). The lowest pH that *B. burgdorferi* is expected to experience in the tick midgut is ∼6.8 after a blood meal (61). This pH is permissive to *B. burgdorferi* proliferation in vitro to saturating densities of >10^8^ cells/mL (60, 79). On the other hand, our in vitro experiments showed that starvation alone (without medium acidification) causes a loss of cell proliferative potential within days, as shown when *B. burgdorferi* cells were resuspended in RPMI 1640 medium (Fig. 3D). While it is suggested that the midgut of unfed ticks is poor in nutrients (10, 11, 14, 15, 21–23), genetic and transcriptomic studies suggest that *B. burgdorferi* may use alternate carbohydrate source (e.g., glycerol, *N*-acetylglucosamine, chitobiose, maltose, mannose, and trehalose) and residual mammalian host blood components available during the tick phase to maintain limited metabolic activity (22, 80–89).

It is also worth noting that the apparent growth arrest of *B. burgdorferi* in unfed ticks is an inference from population observations showing that the spirochete cell count does not significantly change over time (Fig. 5) (16). We propose that the relatively constant spirochete count may reflect a balance between cell growth and death rather than cessation of bacterial replication. Non-sporulating *B. subtilis* cells have been shown to survive extended period (>100 days) of carbon starvation by relying on scarce nutrients released from cell lysis, allowing them to maintain very slow growth (∼4-day doubling time) (78). Similar principles of scavenging biomass from lysed cells have also been suggested to be at play for starving *E. coli* cultures (90). An analogous scenario of extreme nutrient scavenging and slow oligotrophic growth might occur for *B. burgdorferi* in the midgut of unfed ticks. The tick immune system, which has been shown to limit the burden of *B. burgdorferi* within the vector (91), could also contribute to keeping the spirochetal cell density nearly constant. We hypothesize that the maintenance of a slow growth rate of *B. burgdorferi* inside the unfed tick (balanced with cell death) will allow cells to maintain one or more copies of their full genomic complement and thereby retain their proliferative ability until the next blood meal.

Our findings have several practical implications for the *Borrelia* field. First, they invite caution when interpreting results from PI staining. While PI-negative staining indicates membrane integrity, it does not necessarily reflect the ability of cells to proliferate in fresh medium (Figs. 1A and S1F). This is in line with studies on other bacteria, which have shown that cells can lose their reproductive potential without losing their membrane integrity and vice versa (92, 93). Second, the observed loss of plasmids in stationary phase may also contribute to the well-known reported phenomenon of plasmid loss in culture, particularly in high-passaged strains (65–70). This common loss of plasmids has plagued the *Borrelia* field because many plasmids that are not essential for *B. burgdorferi* growth in cultures are required for infectivity (65–70, 94). While we observed rare instances of plasmid loss in exponential phase, the probability of plasmid loss was higher in stationary phase (Fig. 4F). Our data suggest that passaging cultures exclusively in exponential phase may mitigate this long-standing problem of plasmid loss. Third, the rapid loss in genome copies per cell and the ensuing loss of cell proliferative capacities during stationary phase (Figs. 1A, 4, and S5) may also explain anecdotal reports of lower yield and poorer quality of electrocompetent cell and genomic DNA preparations from stationary phase cultures. Our observations strengthen the recommendation made by published protocols to use cultures at density below 10^8^ cell/mL (i.e., before they reach stationary phase) for genetic manipulations (95–99).

Overall, our CFU assays on stationary phase or RMPI 1640 cultures reveal that these in vitro conditions poorly mimic the unfed tick conditions with respect to spirochete persistence. The loss of genome copies in stationary phase cells results in a progressive decrease in genome concentration. Whether such genome dilution occurs in vivo (e.g., in starved ticks or some infected mammalian tissues) will require further experimentation. Though technically challenging, we believe that such experiments may yield interesting hypotheses because genome dilution alone (i.e., without changes in nutrient availability and other environmental conditions) has recently been shown to modulate transcriptome and proteome composition across organisms as diverse as *E. coli*, yeast, and mammalian cells (100–102).

## MATERIALS AND METHODS

### Bacterial strains and general growth conditions

The *B. burgdorferi* strains used in this study are listed in Table S1. These strains were grown in complete Barbour-Stoenner-Kelly (BSK)-II liquid medium at 34°C under 5% CO_2_ atmosphere (40, 103, 104), unless otherwise specified. Complete BSK-II medium contained 50 g/L bovine serum albumin (Millipore #810036), 9.7 g/L CMRL-1066 (US Biological #C5900-01), 5 g/L neopeptone (Difco #211681), 2 g/L yeastolate (Difco #255772), 6 g/L HEPES (Millipore #391338), 5 g/L glucose (Sigma-Aldrich #G7021), 2.2 g/L sodium bicarbonate (Sigma-Aldrich #S5761), 0.8 g/L sodium pyruvate (Sigma-Aldrich #P5280), 0.7 g/L sodium citrate (Fisher Scientific #BP327), 0.4 g/L *N*-acetylglucosamine (Sigma-Aldrich, #A3286), and 60 mL/L heat-inactivated (inactivated at 50°C for 30 min) rabbit serum (Gibco #16120). The pH was adjusted to 7.6 using sodium hydroxide. For testing the effects of medium acidification without nutrient limitation, complete BSK-II medium was prepared, and its pH was adjusted to 6 using L(+)-lactic acid, 90% solution in water (Thermo Scientific #189872500). Tubes contained 6 mL or 14 mL culture volumes depending on the tube size used (8-mL volume, Falcon, #352027, or 16-mL volume, Falcon, #352025) and were kept tightly closed. Any larger volume vessels were kept loosely capped in the incubator. The pH of the culture medium was measured with a pH meter following filtering the culture through a 0.1-μm filter (Millipore # SLVVR33RS).

### Growth curve generation

The growth of *B. burgdorferi* cultures was monitored by determining cell density expressed as cells/mL. Cell density was measured by using a Petroff-Hausser chamber (C-Chip disposable hemocytometer by INCYTO). The culture was diluted in 1X phosphate-buffered saline (PBS), and a 10 μL sample was loaded for counting. Cells were directly counted under darkfield illumination using a Nikon Eclipse E600 microscope equipped with a 40×, 0.55 numerical aperture (NA) Ph2 phase-contrast air objective and darkfield condenser optics.

*B. burgdorferi* cultures were also evaluated for their ability to form colonies using a semisolid BSK-agarose plating method (95, 98). Briefly, the cultures were serially diluted 10-fold in fresh BSK-II medium, and a ten-centimeter Petri dish was seeded with 0.5 or 1 mL of each serial dilution. After a brief (≤ 5 min) pre-equilibration at 55°C, three parts BSK-1.5 medium (containing 69.4 g/L bovine serum albumin, 12.7 g/L CMRL-1066, 6.9 g/L neopeptone, 3.5 g/L yeastolate, 8.3 g/L HEPES, 6.9 g/L glucose, 6.4 g/L sodium bicarbonate, 1.1 g/L sodium pyruvate, 1.0 g/L sodium citrate, 0.6 g/L *N*-acetylglucosamine, and 40 mL/L heat-inactivated rabbit serum) was mixed with two parts of sterile, 55°C-equilibrated, 1.7% agarose solution in water to generate a plating mixture. Approximately 25 mL of the plating mixture was poured onto each pre-seeded plate, and plates were gently swirled to mix before allowing them to solidify at room temperature. Once solidified, the plates were transferred to a humidified 5% CO_2_ incubator and incubated between ten days and three weeks until visible colonies appeared. Then, the colonies on each plate were counted and the culture density was subsequently calculated and reported as colony forming units per mL (CFU/mL).

*B. burgdorferi*’s ability to grow in culture was also measured using a limiting dilution method, adapted from prior studies (106). Briefly, 10-fold serial dilutions of a culture were made using fresh BSK-II medium. For each dilution, eight wells of a 96-well plate were seeded with 250 μL of each respective serial dilution per well. The plates were incubated for up to two weeks in a humidified, 5% CO_2_ incubator, after which each well was scored for growth. Wells that changed color from pink to orange-yellow due to growth-dependent acidification of the medium were considered positive. Wells that remained pink were inspected for spirochete growth using a darkfield microscope. The concentration of growth-capable cells in the parental culture was then calculated using the method of Reed and Muench (107) and expressed as tissue culture infectious dose 50 per mL (TCID_50_/mL).

### Cell sampling from cultures

Large volume cultures (∼70-130 mL) of strains B31-MI, 297, and N40 were grown in complete BSK-II medium at 34°C under 5% CO_2_ atmosphere. At each time point, a small aliquot of culture (∼5-10 mL) was taken and briefly vortexed before being characterized by counting cell culture density, plating for CFU determination, and evaluating PI uptake and Hoechst 33342 staining. Entrance into stationary phase was defined as the day a culture reached ∼10^8^ cells/mL (typical range ≥ 9.75x10^7^ – 2.5x10^8^ cells/mL for the onset of stationary phase). This day within the time course was designated as day 0, and all other time course data collection points were standardized relative to the start of stationary phase.

### Starvation in RPMI 1640 medium

Cultures of strains K2, Bb914, and N40 were grown to densities of no more than 5x10^7^ cells/mL in complete BSK-II medium. A small volume of culture was set aside for imaging PI uptake as a pre-resuspension in RPMI 1640 condition. The remaining cells were pelleted using a 4,300 × g spin for 10 min in an Allegra X-14R centrifuge (Beckman Coulter) equipped with a swinging bucket SX4750 rotor. Cells were then resuspended in an equivalent volume of RPMI 1640 Medium ATCC Modification (Gibco #A1049101). The cultures were subsequently tracked for cell culture density, colony-forming ability, and PI uptake for the duration of the experiment.

To generate the pH curve of K2 cells during RPMI incubation, two cultures of strain K2 were grown to 10^6^ cells/mL, then subsequently pelleted, resuspended, and incubated in the same volume of RPMI 1640 medium at 34°C. The culture pH was measured as described above.

### Culture in acidic BSK-II medium

Cultures of strain K2 were grown to densities below 5x10^7^ cells/mL in complete BSK-II medium. A small volume of each culture was set aside for determining cell density, colony-forming abilities, and PI uptake before resuspension in acidic medium; this time point served as day 0 in the experiment. The remaining cells were pelleted using a 4300 × g spin for 10 min in an Allegra X-14R centrifuge (Beckman Coulter) equipped with a swinging bucket SX4750 rotor and then resuspended in an equivalent volume of complete BSK-II medium adjusted to pH 6.0 as described above. Culture density was determined by darkfield counting and serial dilutions were plated in semisolid BSK-agarose for CFU measurements for all subsequent experiment days. Aliquots of cultures were also imaged by fluorescence microscopy for PI uptake.

### Culture at room temperature

Two 7-mL cultures of *B. burgdorferi* strain CJW_Bb379 were inoculated at 10^4^ cells/mL in BSK-II medium and incubated at room temperature (∼21°C) in the dark in closed tubes. Culture density was determined by darkfield counting and serial dilutions were plated in semisolid BSK-agarose for CFU measurements. Aliquots were imaged by fluorescence microscopy.

### Microscopy

For fluorescence imaging, *B. burgdorferi* strains were spotted onto a 2% agarose-PBS pad, as described previously (64, 108), covered with a no. 1.5 coverslip, then imaged using Nikon Eclipse Ti microscopes equipped with either a 100X Plan Apo 1.45 or 1.40 NA phase-contrast oil objective, a Hamamatsu Orca-Flash4.0 V2 CMOS camera, and either a Sola Light Engine (Lumencor) or pE-4000 (CoolLED) light source. The microscopes were controlled by the Nikon Elements software. The following Chroma filter cubes were used to acquire the fluorescence images: DAPI, excitation ET395/25x, dichroic T425lpxr, emission ET460/50 m; GFP: excitation ET470/40x, dichroic T495lpxr, emission ET525/50 m; mCherry/TexasRed, excitation ET560/40x, dichroic T585lpxr, emission ET630/75 m. DNA staining was obtained by incubating the culture for at least 15 min with Hoechst 33342 (Molecular Probes #H3570) at a final concentration of 2 μg/mL, and membrane integrity was assessed by incubating culture samples for at least 15 min with propidium iodide (Invitrogen #P3566) at a final concentration of 1 μg/mL. Both dyes were directly added to the culture sample, and no washes were performed.

### Image analysis

Unless otherwise specified (i.e., cases where cell outlines were not produced), cell outlines were generated from phase-contrast images using the Oufti software package and used for subsequent image analysis, with the following parameters: Edgemode, LOG; Dilate, 2; openNum, 3; InvertImage, 0; ThreshFactorM, 0.985; ThreshMinLevel, 0; EdgeSigmaL, 1; LogThresh, 0. Cell outlines were manually curated to remove outlines of cell debris and outlines of cells that curled on themselves, intersected with other cells, or extended beyond the field of view. In some cases, outlines of cells were manually joined to cover the full length of the cell. For downstream analyses of Oufti-generated cell outlines, cells with lengths < 5 μm were removed, as these corresponded to debris that were erroneously assigned as cell meshes.

To quantify the percentage of PI-negative cells for B31-MI, 297, N40, and K2 in pH 6.0 BSK-II medium, fluorescence images in the TexasRed channel were background-subtracted. Data were analyzed using the custom MATLAB script analyze_signal_intensity_Oufti.m (written for this study). To determine whether cells were PI-positive or -negative, the first and last days of the time course for each respective strain of one replicate were compared on a histogram of mean cell intensities. Since these signals diverged into a bimodal distribution, an Otsu threshold (110) was applied to find the split between the two states. An Otsu threshold was determined for each strain, as the bimodal split varied between strains except for strain K2 in pH 6.0 BSK-II medium; in that case, most cells were PI-negative by distribution, and an accurate Otsu could not be determined. Therefore, the Otsu threshold used for B31-MI was applied to K2 since those strains are closely related (111). Cells for which the fluorescent PI signal was greater than the Otsu threshold were deemed “PI-positive” while those below the Otsu threshold were considered “PI-negative”. To determine the number of DNA signal objects in a cell, the objectDetection function of Oufti (109) was used on manually curated cell outlines. The following parameters were used: manual background threshold, 0.1; background subtraction method, 4; background subtraction threshold, 0.1; background filter size, 8; smoothing range (pixels), 3; magnitude of LOG filter, 0.1; sigma of PSF, 1.62; fraction of object in cell, 0.1; minimum object area, 10. The number of detected nucleoid objects was automatically exported from the analyzed cells after using objectDetection when possible using the MATLAB function get_nuc_num.m; The percentage of cells with one object detected in the analyzed population of cells was plotted.

Hoechst intensity data, presented in Fig. S4E-G, was measured from the same custom MATLAB script analyze_signal_intensity_Oufti.m. Mean cell intensities were concatenated in a large data structure containing all strains, replicates, and days of measurement. The mean intensity through culture age was fitted to an exponential decay function (Eq. 1 below). The terms of the equation yield three parameters: the initial mean fluorescence value *y*_0_, the rate constant λ, and an arbitrary constant *C* for determining the basal fluorescence state of the fit. All parameters were left floating for the fit with no boundaries. The half-life of DNA stain in cells was inferred by 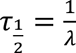.

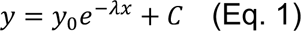

Fluorescent puncta detection for *oriC* and/or plasmid copy number determination were performed as previously described (52). Briefly, the MATLAB script Modified_Find_Irregular_Spots.m was used, with the parameters: fitRadius: 5, edgeDist: 2.5, centerDist: 1; peakRadius: 3; shellThickness: 1; quantileThreshold: 0.3. For each microscopy experiment and fluorescence channel, the intensityRatioThreshold parameter was determined by empirical testing and visual inspection (see data available as supplemental material for reporting of intensityRatioThreshold parameters). Cell outlines were visually inspected for detection of fluorescent puncta. Cell outlines that had clear *oriC* foci, but an incorrectly detected number of spots were removed from all subsequent analyses. For quantification of *oriC* foci, cell outlines without clear *oriC* foci were manually counted and tabulated for inclusion in the final analysis. These excluded cells (most often displayed for strains CJW_Bb378 and CJW_Bb379 grown at 34°C in BSK-II during stationary phase; see Figs. 4B and S4C-D) often had very low fluorescence signal or patchy fluorescence signal, leading to a frequent misassignment of spots in subsequent spot detection MATLAB functions. From this manually curated cell list, *oriC* spots were identified and added to the cell list using add_spots_to_cellList.m, and the final data were exported into a table format using export_to_table.m and extract_field.m (52). Cells identified as having “NaN” or undefined values for spots were removed from the final table. The manually counted and tabulated cells without clear *oriC* foci were then included in the final analyses for determining *oriC* per cell and the percentage of cells in the population without *oriC* foci.

For images from strains in which both *oriC* and an endogenous plasmid were labeled (see Table S1 for details), the following steps were taken. First, any cells without fluorescence signal were removed; these cells likely had lost the shuttle vector used to label both *oriC* and a given endogenous plasmid (52). Next, cells were analyzed for *oriC* puncta. As before, cell outlines that had clear *oriC* foci, but an incorrectly detected number of spots were removed from all subsequent analyses. Then, cells without clear *oriC* spots were also removed but again manually tracked and counted; these cells were removed since these cells are unlikely to be able to proliferate without a full intact copy of the chromosome. From this dataset, cells were manually curated for spot detection and *oriC* and plasmid spots were added to the cell list using add_spots_to_cellList. The final data were exported into a table format using export_to_table.m and extract_field.m (52). The fluorescence intensity profiles for cells of strain CJW_Bb489 were generated as previously described (109).

For analysis of plasmid retention and plasmid retention variance between exponential and stationary phase cultures, the relative numbers of plasmid copies per cell were compiled in a large Pandas (112, 113) data frame via Python using its class function *groupby*. The data frame was grouped by plasmid (genetic element), growth phase (exponential vs. stationary), and replicate (1 vs. 2). Mean plasmid spots for each replicate were determined first, then the standard deviation of each genetic element was applied to yield Fig. 4G. The fraction of cells without plasmids in Fig. 4F were determined through a separate *groupby* function by counting the number of observed cells without plasmid foci, then dividing that value by the total number of cells for each replicate. The mean fraction of cells without plasmids was determined for each replicate.

For experiments related to starvation in RPMI 1640, cell outlines were not generated due to the high frequency of cells with non-spirochetal morphologies; instead, cells were manually counted to distinguish between cells with the normal flat-wave shape and cells with various morphological defects that included membrane bulges and condensed ball-like forms (i.e., round bodies). Cell counting was facilitated by the Fiji cell counter plugin cell_counter.jar. The percentage of round body cells in the population was subsequently determined.

### Calculation of propidium iodide-negative cells

To estimate the total number of PI-negative cells in a population, the percentage of PI-negative cells determined by microscopy was multiplied by the number of cells counted using the Petroff-Hausser chamber. The subsequent total number of PI-negative cells was reported as cells/mL. The raw percentages of PI-negative cells in a population were also plotted on a linear scale and provided in supplementary figures.

### Plasmid profiling

The endogenous plasmid complement of B31-derived clones was determined by multiplex PCR, using previously described primers (71) and the DreamTaq DNA polymerase Green PCR Master Mix kit (ThermoFisher). To monitor plasmid loss, two cultures of CJW_Bb523 were grown in parallel. These cultures were then plated in exponential phase, after five days in stationary phase, and after ten days in stationary phase, all without any passaging steps. Colonies were transferred to in 1.5 mL BSK-II medium at 34°C under 5% CO_2_ atmosphere. After growth, cells were harvested by pelleting at 10.000 × g spin for ten minutes using an Eppendorf 5430R centrifuge equipped with a FA-45-30-11 rotor. The resulting pellet was resuspended in 50-100 μL water. This suspension was directly used as source of DNA in the multiplex PCR reactions. Eleven clones were isolated and tested per culture in exponential phase, while twelve clones were isolated and tested per culture per time point in stationary phase.

### Tick studies

In a recent study (52), larval *Ixodes scapularis* ticks were colonized with *B. burgdorferi* strain CJW_Bb474 by feeding on CJW_Bb474-infected 4-8 weeks-old female RML (*Mus musculus*) mice, an outbred strain of Swiss-Webster mice reared at the Rocky Mountain Laboratories breeding facility. These tick larvae were maintained in the laboratory at ambient light and temperature in bell jars over potassium sulfate-saturated water. The larvae molted into nymphs, which were then kept in the same conditions until they were mechanically homogenized in BSK-II medium. The resulting suspensions were plated in semisolid BSK-agarose medium, which over time yielded *B. burgdorferi* colonies that were counted. These colony counts were used to calculate the viable spirochete load in each unfed nymph. Six unfed nymphs were plated approximately two months after their larval feeding (approximately one month after their molt) and the result was reported in the previous study (52). An additional five unfed nymphs were plated approximately 15 months after their larval feeding, or 14 months after their molt, and are reported here for the first time. For each of these five nymphs, between eight and ten *B. burgdorferi* clones per nymph were grown in BSK-II medium and their plasmid content was determined by multiplex PCR as mentioned above.

### Statistical tests

For Fig. 4F-G, an unpaired *t* test with Welch’s correction was performed in GraphPad Prism to compare between exponential phase and two days in stationary phase conditions. For Fig. 5B, the log-normal CFU data were converted to a linear, normal scale via a log10 transformation, then were compared using an unpaired *t-*test with Welch’s correction in GraphPad Prism version 10.2.2 software.

### Data visualization

All data were plotted to generate figures using GraphPad Prism version 10.2.2 software, with the exception of the plots in Fig. S4E-G, which were done using MATLAB. Images were visualized using Fiji (114) and Nikon NIS Elements. Background from fluorescence images was subtracted in Fiji by measuring the mean intensity of a region of interest box drawn on a portion of the image where there were no cells. This determined mean intensity of pixels was used to subtract the value from all pixels within the image. Images were additionally adjusted in Fiji, as follows: to display PI-staining intensities in Fig. 3B, background from fluorescence images was subtracted, images were scaled within the same intensity values, and the Red Hot look up table was applied. For fluorescent images in Figs. 4A, 4E, S4A, and S5E, the mean background was subtracted from the raw fluorescence images in Fiji, and image pixel intensity values were autoscaled for display. For fluorescent images in Fig. 4D, the mean background was subtracted from the raw fluorescence images in Fiji, and image pixel intensity values were scaled to be the same between exponential and stationary phase images within a given strain (CJW_Bb274 and CJW_Bb517) and fluorescent marker (mCherry for *oriC* and GFP for plasmid). For all phase contrast images displayed (Figs. 1B, 3B, 4A, 4E, S4A, S4A and S5E), image pixel intensity values were autoscaled for display. All images were converted to RGB format for display. Adobe Illustrator 2024 was used to generate figures.

## DATA AVAILABILITY

Raw images (.nd2 file format) acquired and analyzed for this study will be available on the public Biostudies repository. Reporting of specific exposure times and camera information for microscopy experiments, as well as specific n-values and intensityRatioThreshold parameters used during subsequent image analysis, are provided as Supplemental File 1. All source data used to generate figures are provided as Supplemental File 2.

## CODE AVAILABILITY

All code developed as part of this study is publicly available on Github at https://github.com/JacobsWagnerLab/published/tree/master/Zhang_Takacs_et_al_2024.

## ACKNOWLEDGMENTS

We would like to thank the Jacobs-Wagner laboratory for support, discussion, and feedback on the manuscript. J. Z was in part supported by the Stanford predoctoral training grant in Cellular and Molecular Biology (T32GM007276). C.N.T. was supported in part by an American Heart Association postdoctoral fellowship (18POST33990330). P.A.R. and J.W. were supported by the Intramural Research Program of the National Institute of Allergy and Infectious Diseases, National Institutes of Health. C.J.-W. is an investigator of the Howard Hughes Medical Institute.

## AUTHOR CONTRIBUTIONS

J.Z., C.N.T., E.A.M., and C.J.-W. designed the research. J.W. and P.A.R. performed the tick-mouse transmission studies and plating of crushed ticks. J.Z., C.N.T., and E.A.M. collected the data for the studies. J.Z., C.N.T., and J.B. curated and analyzed images. J.W.M. and Y.T. developed code used for analysis, with input from J.Z. and C.J.-W. J.W.M. assisted with statistical analyses performed throughout the paper. J.Z. and C.N.T. made the figures with input from C.J.-W. J.Z. and C.J.-W. wrote the paper with input from C.N.T. and the other authors. C.J.-W. obtained funding and oversaw the project.

---

**Figure S1.**
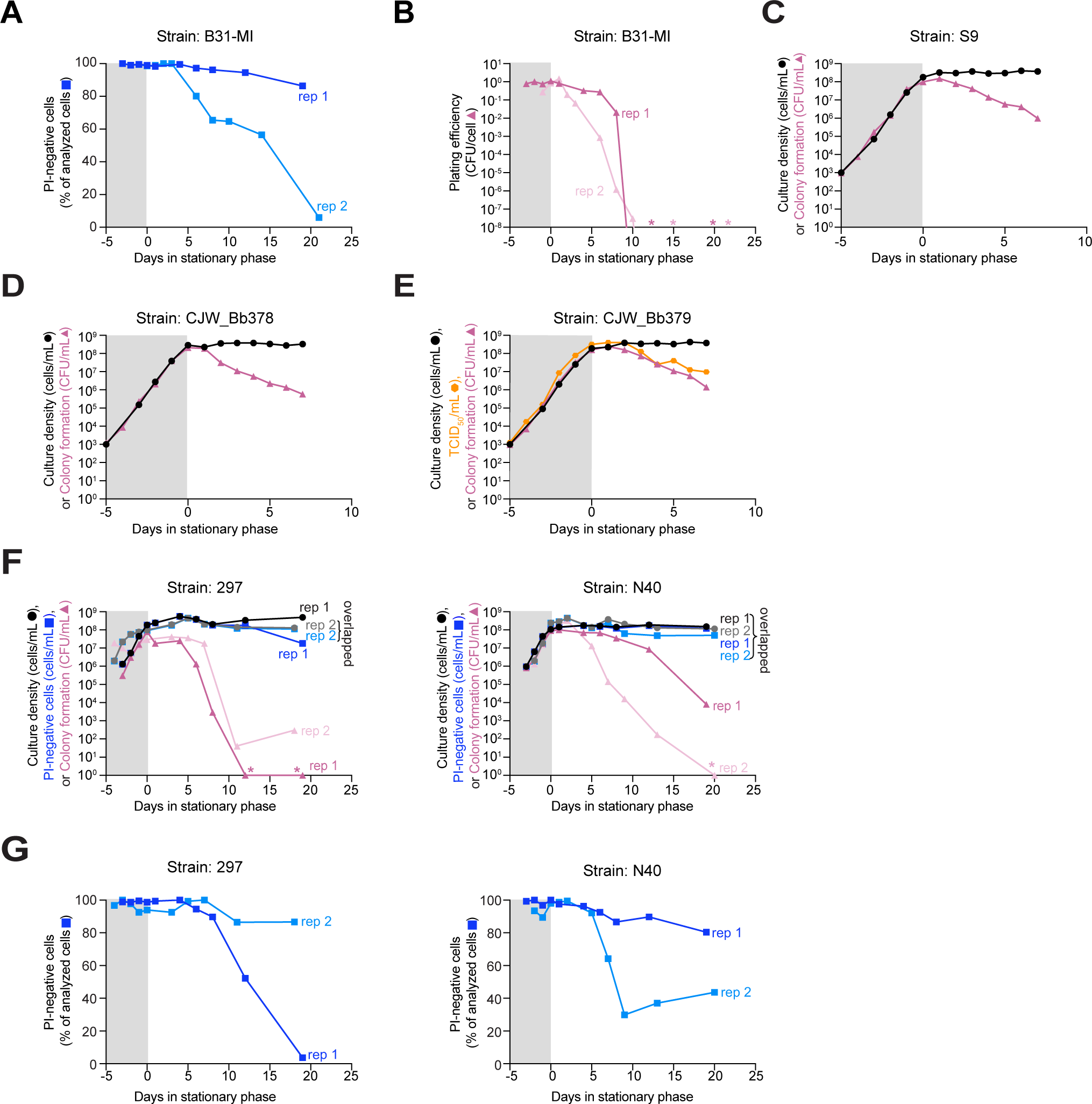
Stationary phase phenotypes characterized in various strains. Gray and white backgrounds indicate exponential and stationary phases, respectively. Plot showing the percentage of cells (n = 72-417 for each strain and time points; see Supplemental File 1 for specific n values) from cultures used in Fig. 1A that remained negative for propidium iodide (PI) staining shown on a linear scale; these values were used to calculate the number of PI-negative cells. Results from two independent cultures (biological replicates rep 1 and 2, represented by dark and light blue squares) of strain B31-MI are shown. **A.** Plot showing the plating efficiency for B31-MI. Plating efficiency was calculated by dividing the CFU/mL of a culture by the number of cells/mL measured at each time point (shown in Fig. 1A). Asterisks indicate that the plating efficiency could not be plotted on a log scale, as the value was zero. Results from two independent cultures (biological replicates rep 1 and 2, represented dark and light pink triangles) of strain B31-MI are shown. **B.** Plot showing culture densities (cells/mL, black circles) in comparison to colony-forming ability (CFU/mL, pink triangles) for a single culture of the clonal B31-derived strain S9. **C.** Same as in (C) for clonal B31-derived strain CJW_Bb378. **D.** Same as in (C) but for clonal B31-derived strain CJW_Bb379. Here, the density of viable cells in the culture was also determined by a liquid culture using microtiter plate-based limiting dilution assay. This density of viable cells is expressed as tissue culture infectious dose 50 (TCID_50_) per mL (shown as orange hexagons). **E.** Same as in Fig. 1A but for two independent cultures (biological replicates, rep 1 and 2) of strains 297 (left) and N40 (right). Black and grey circles represent culture density, dark and light blue squares represent PI-negative cells, and dark and light pink triangles represent CFU/mL for rep 1 and rep 2, respectively. Asterisks indicate that no colonies were detected. **F.** Plots showing the percentage of PI-negative cells in the same cultures as used in (F) shown on a linear scale.

**Figure S2.**
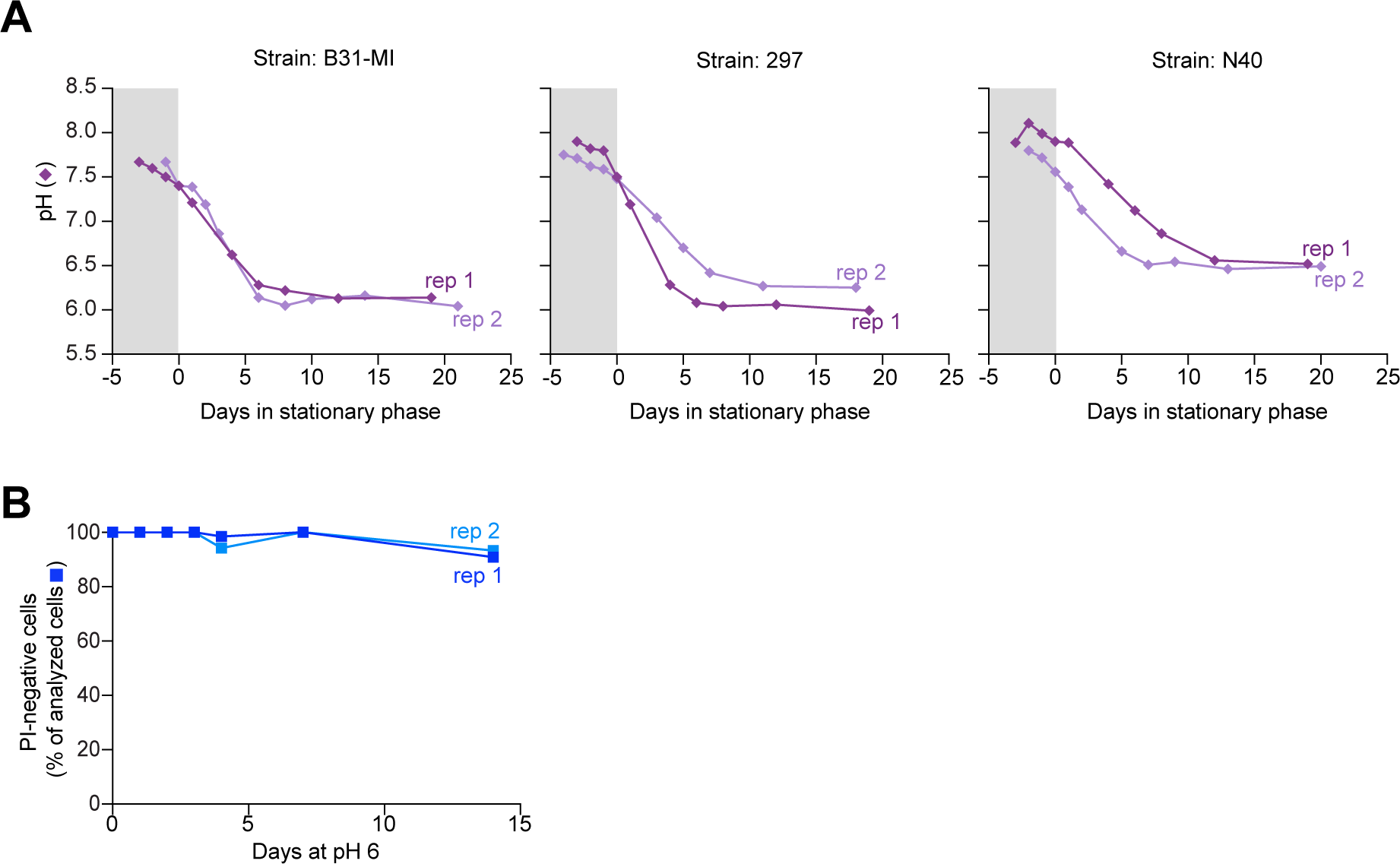
Medium acidification in cultures and its effect on membrane permeability. **A.** Plots showing the changes in pH that occurred during cultivation of strains B31-MI, N40, and 297 in BSK-II medium. These pH measurements were done using the same cultures as those for Figs. 1 and S1F-G. Gray and white backgrounds indicate exponential and stationary phases, respectively. Shown are results from two independent cultures (biological replicates, rep 1 and 2) of each indicated strain. **B.** Plot showing the percentage of K2 cells that remained negative for propidium iodide (PI) uptake after culture at pH 6.0. There are the same results as in Fig. 2B, except on a linear scale.

**Figure S3.**
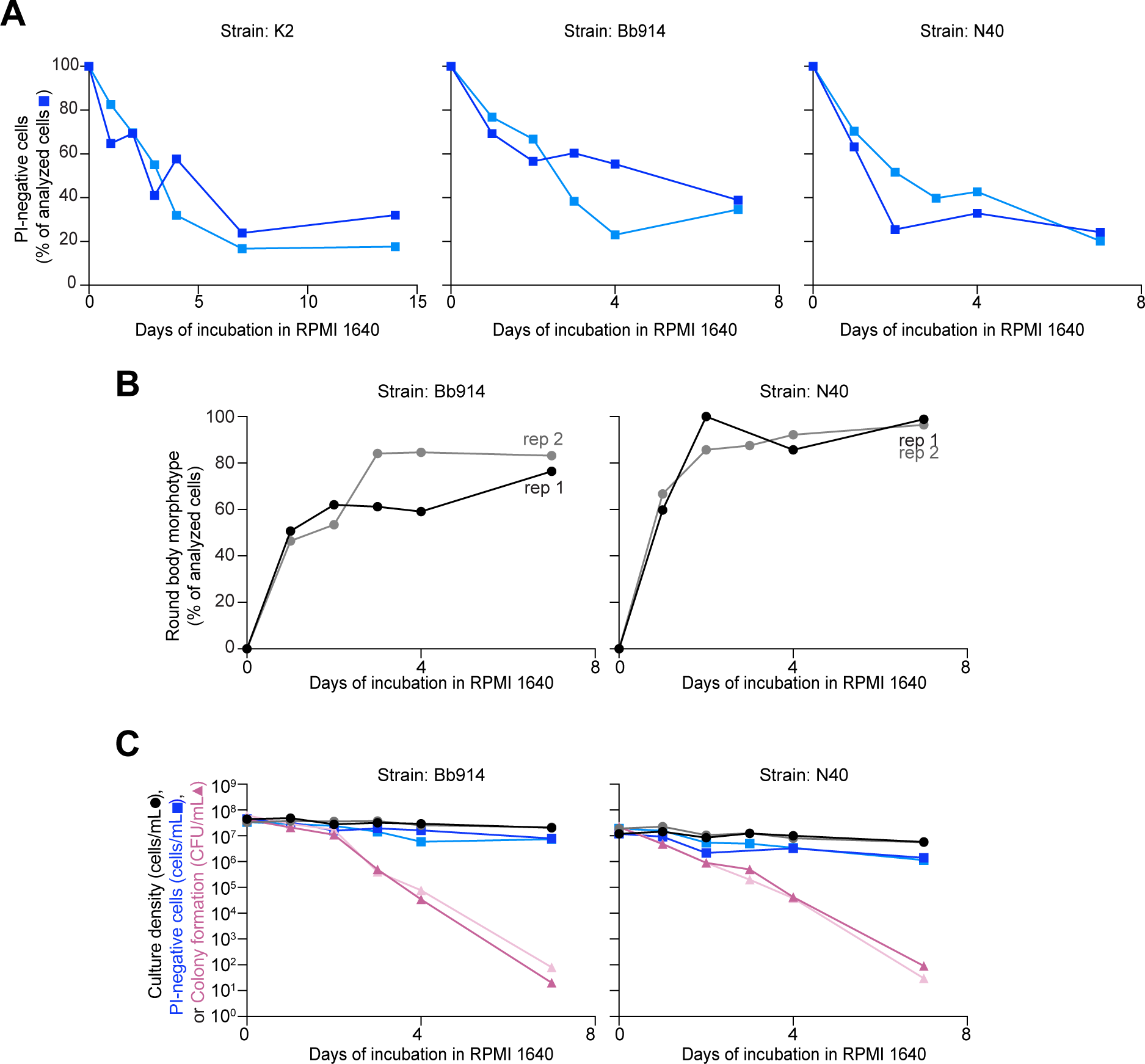
Effects of starvation in RPMI 1640 on various *B. burgdorferi* strains. Results from two independent cultures (biological replicates, rep 1 and 2) of each respective strain are shown. See Supplemental File 1 for specific n values for each time point and strain. **A.** Plots showing the percentage of cells from cultures of strain K2 (as used in Fig. 3), strain Bb914 (a derivative of strain 297), and non-clonal strain N40 that remained negative for propidium iodide (PI) uptake shown on a linear scale. For PI-negative cell determination, 48 to 311 cells were analyzed for each strain and time point. **B.** Same as Fig. 3C for cultures of strains Bb914 and N40. For round-body determinations, 70 to 311 cells were analyzed for each strain and time point. **C.** Same as Fig. 3D for cultures of Bb914 and N40. For PI-negative cell determination, 70 to 311 cells were analyzed for each strain and time point.

**Figure S4.**
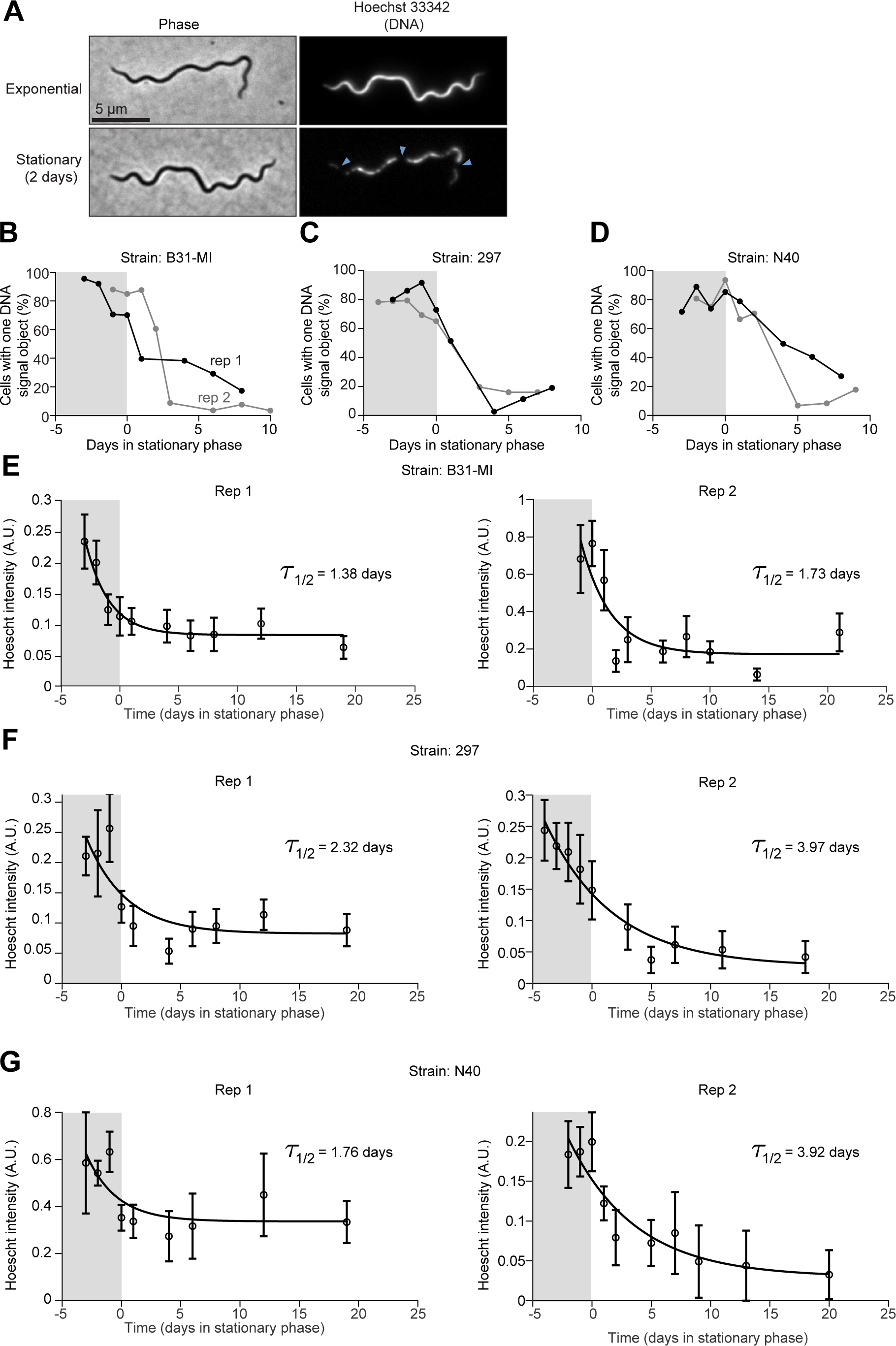
Changes to DNA staining patterns in stationary phase cells. For panels B-G, gray and white backgrounds indicate exponential and stationary phases, respectively. Results from two independent cultures (biological replicates, rep 1 and 2) of each respective strain are shown. See Supplemental File 1 for specific n values for each time point and strain. **A.** Representative images of a cell from an exponential or stationary phase culture of B31-MI where DNA was visualized by staining with Hoechst 33342. Light blue arrowheads indicate gaps depleted of DNA signal in the stationary phase cell. **B.** Plot showing the percentage of the cell populations with one continuous DNA signal for the B31-MI cultures used in Fig. 1. For DNA object detection analysis, 72 to 418 cells were analyzed for each strain and time point. **C.** Same as in (B) except for cultures of strain 297, which were the same cultures as those used for Fig. S1F. For DNA object detection analysis, 66 to 332 cells were analyzed for each strain and time point. **D.** Same as in (B) except for cultures of strain N40, which were the same cultures as those used for Fig. S1F. For DNA object detection analysis, 45 to 350 cells were analyzed for each strain and time point. **E.** Plot showing the decay of the Hoechst signal intensity in BM31-MI cells as a function of culture age. Mean whole cell intensity of either biological replicate for each time point was fit to an exponential decay to illustrate the sharp decrease in signal. The same B31-MI culture was used as in Fig. 1. A.U. indicates arbitrary units. **F.** Same as in (E) except for cultures of strain 297. **G.** Same as in (E) except for cultures of strain N40.

**Figure S5.**
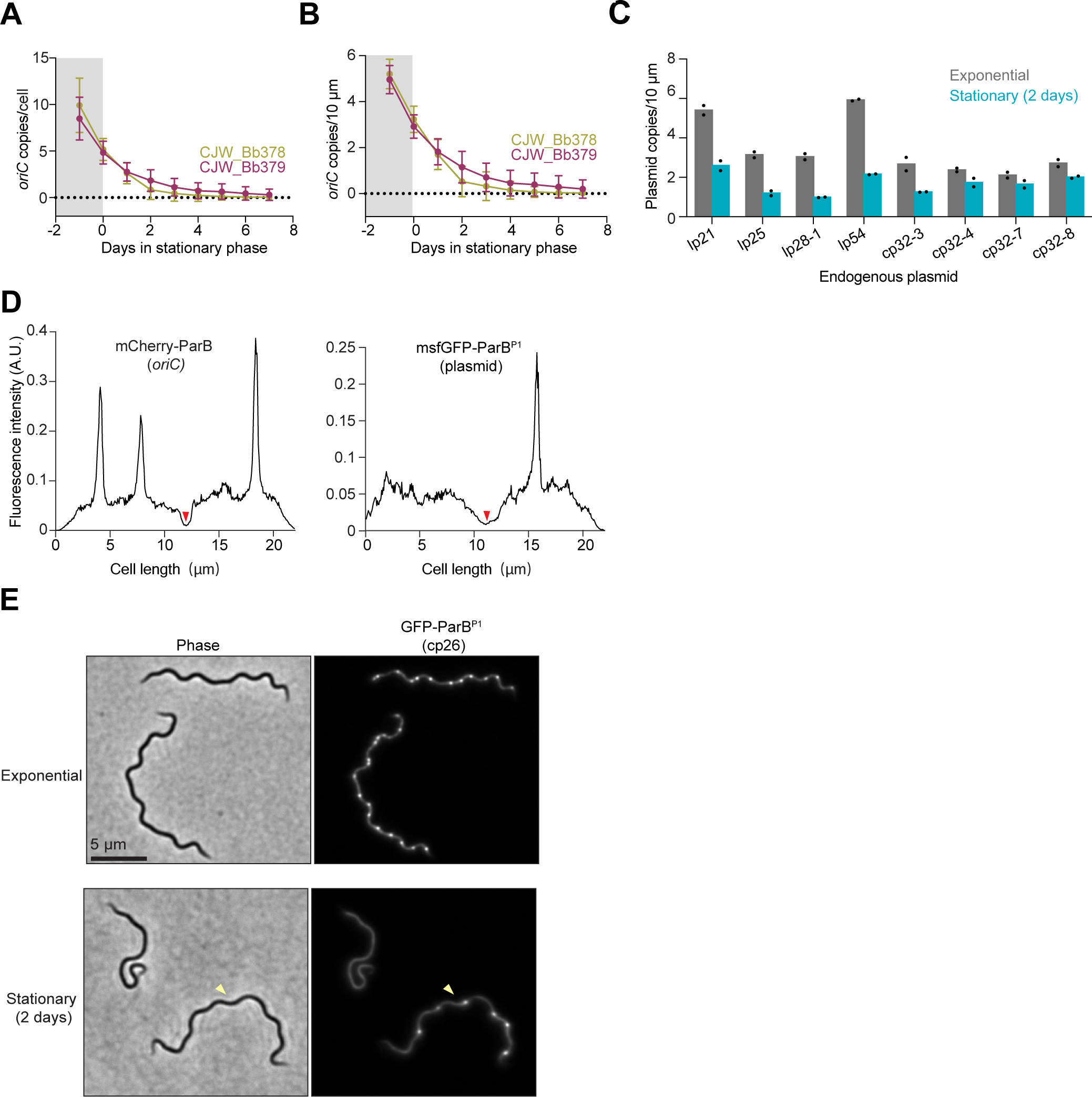
Decrease in *oriC* and plasmid copy density during stationary phase. **A.** Plot showing changes in *oriC* copies per cell in cultures of strains CJW_Bb378 and CJW_Bb379 (both clonal B31 derivatives) in BSK-II medium. The numbers of *oriC* copies were determined from fluorescence microscopy images by counting the fluorescent foci of mCherry-ParB (CJW_Bb379) or ParZ-GFP (CJW_Bb378) that indicate the subcellular location of the endogenous *parAZBS* region adjacent to *oriC* (52). One culture for each strain was analyzed at the indicated time points. Shown are means ± standard deviations across cells. For each strain and time point, 60 to 387 cells were analyzed (see Supplemental File 1 for specific n values). Gray and white backgrounds indicate exponential and stationary phases, respectively. **B.** Same as in (A) except that the mean *oriC* densities (expressed as *oriC* copies per 10 μm of cell length) are plotted. **C.** Plot showing plasmid densities (expressed as plasmid copy number per 10 μm of cell) in exponential phase (grey bars) and after two days in stationary phase (teal bars) using the same cultures as in Fig. 4C. Each black dot represents an independent biological replicate. Only cells (n = 22-167) with at least one clear *oriC* focus were considered in this analysis. The strain identities and the number of cells analyzed for each data point are detailed in Supplemental File 1. **D.** Plot showing the intensity profiles for mCherry-ParB and GFP-ParB^P1^ signals along the cell length for the CJW_Bb489 cell shown in Fig. 4E. The division site, reflected by the dip in fluorescence signal, is indicated by red arrowheads. A.U. stands for arbitrary units. **E.** Representative phase contrast and fluorescence images of cells of strain CJW_Bb203 in which cp26 is labeled with msfGFP-ParB^P1^. Cells were from a population in exponential phase or in stationary phase for two days. Yellow arrowheads point to a stationary phase cell with clear fluorescent cp26 foci.

**Figure S6.**
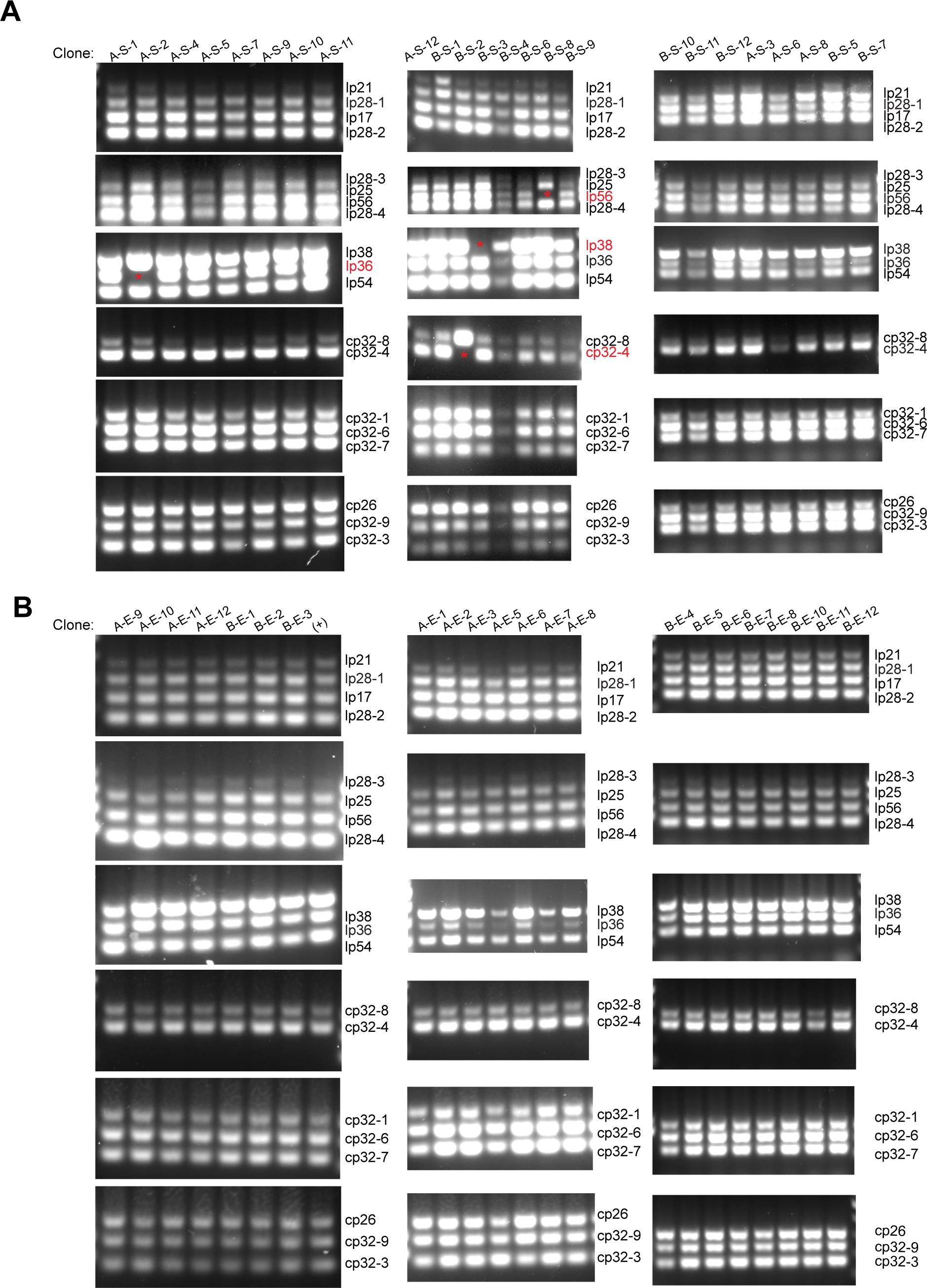
Assessment of the plasmid content in *B. burgdorferi* clones isolated after growth in laboratory cultures. Multiplex PCR was done using plasmid-specific primer pairs previously validated for use on strain B31 (71) and grouped in six sets, as shown in the images. The PCR products were separated by electrophoresis and visualized by SYBR Safe staining and automated detection using a Bio-Rad ChemiDoc Imaging System gel imager. The intensity of the resulting images was scaled to allow for visual detection of the weakly positive bands. As a result of acquisition and scaling, some of the more intense bands are saturated. All multiplex PCR results for all clones analyzed are summarized in Table 1. **A.** Gel images for PCR products obtained by multiplex PCR profiling of all clones tested from 10-day-old stationary phase culture of CJW_Bb523. Lost plasmids (written in red) are indicated with a red asterisk on the gel image. Each clone tested is given a unique identifier: first A or B, corresponding to biological replicate 1 or 2, then S for stationary phase, followed by the identification number of the screened colony (see Table 1 for details). **B.** Same as in (A) except that from cultures in exponential phase. Each clone tested is given a unique identifier as described in (A) except that E is for exponential phase. The (+) indicates a positive control sample, which corresponds to strain CJW_Bb523 isolated in exponential phase.

**Figure S7.**
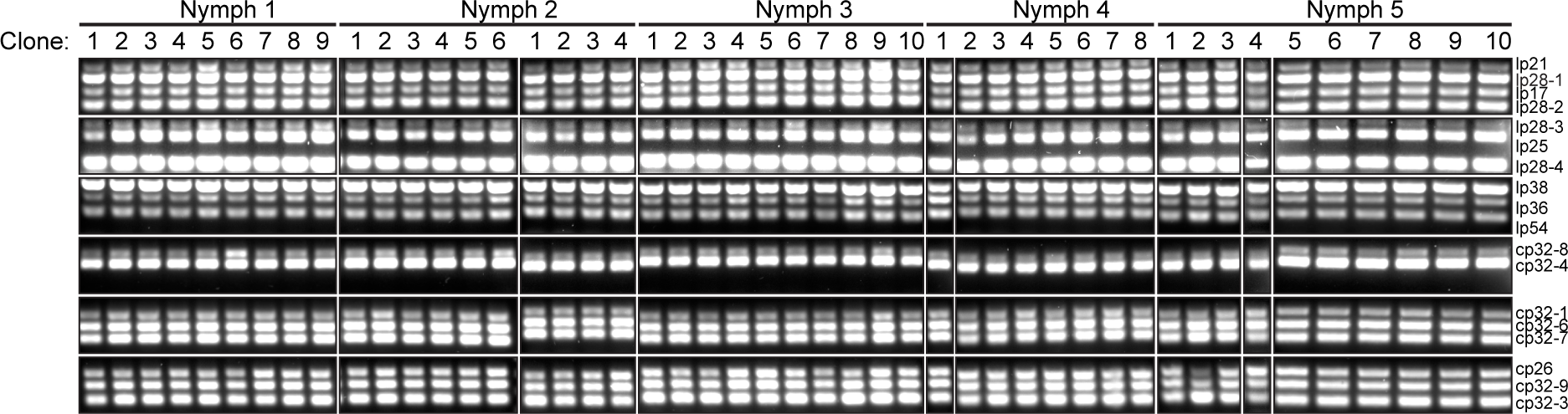
Assessment of the plasmid content in *B. burgdorferi* clones isolated from unfed ticks. Same as in Fig. S6 except that the clones were isolated from the unfed ticks used for Fig. 5. Shown are gel images of PCR products from clones obtained by plating crushed nymphs colonized with strain CJW_Bb474 and maintained unfed at room temperature for 14 months after molt. CJW_Bb474 is a clonal B31-MI derivative that lacks lp5, cp9, and lp56. Isolated and tested clones are grouped by nymph.

**Figure S8.**
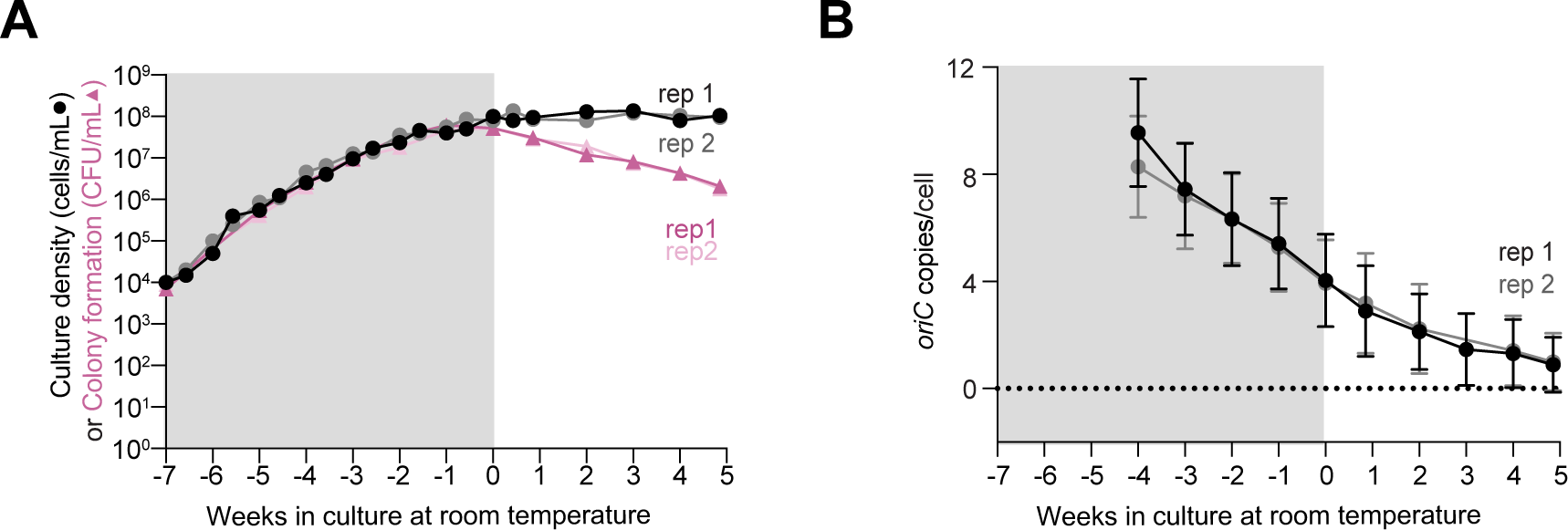
Determination of cell proliferative potential and *oriC* copy number per cell for cultures grown at room temperature. Gray and white backgrounds indicate exponential and stationary phases, respectively. Results from two independent cultures (biological replicates, rep 1 and 2) of strain CJW_Bb379 are shown. See Supplemental File 1 for specific n values for each time point. **A.** Plot showing culture densities (cells/mL, black and gray circles) in comparison to colony forming ability (CFU/mL, dark and light pink triangles) for two cultures of CJW_Bb379 grown in BSK-II at room temperature (∼21°C). Due to the slower growth rate associated with the lower temperature, the x-axis is standardized relative to weeks in stationary phase. **B.** Plot showing changes in *oriC* copies per cell over time for the two cultures of CJW_Bb379 grown in BSK-II at room temperature (∼21°C) using the cultures from (A). For *oriC* copy number quantification, 7 to 811 cells were analyzed for each strain and time point.

**Table S1.** List of strains used in this study.

| Strain | Description | Reference |
| --- | --- | --- |
| B31-MI | Type strain (non-clonal) | (1–3) |
| N40 | Widely used strain (non-clonal) | (4, 5) |
| 297 | Widely used strain (non-clonal) | (6–9) |
| Bb914 | 297 clonal derivative that expresses cytosolic GFP | (10) |
| K2 | Transformable, infectious B31 clone | (11) |
| S9 | Transformable, infectious B31 clone | (11) |
| CJW_Bb203 | S9 derivative with fluorescently labeled cp26 copies and *oriC* copies | (12) |
| CJW_Bb274 | S9 derivative with fluorescently labeled lp25 and *oriC* copies | (12) |
| CJW_Bb326 | S9 derivative with fluorescently labeled lp54 and *oriC* copies | (12) |
| CJW_Bb378 | S9 derivative with *oriC* labeled by ParZ-GFP | (12) |
| CJW_Bb379 | S9 derivative with *oriC* labeled by mCherry-ParB | (12) |
| CJW_Bb474 | CJW_Bb379 derivative that expresses cytosolic GFP | (12) |
| CJW_Bb489 | S9 derivative with fluorescently labeled lp28-1 copies | (12) |
| CJW_Bb515 | B31 derivative with fluorescently labeled cp32-3 copies | (12) |
| CJW_Bb516 | B31 derivative with fluorescently labeled cp32-7 copies | (12) |
| CJW_Bb517 | B31 derivative with fluorescently labeled cp32-4 copies | (12) |
| CJW_Bb518 | B31 derivative with fluorescently labeled cp32-8 copies | (12) |
| CJW_Bb523 | B31 clone, lacks lp5 and cp9 | (12) |
| CJW_Bb526 | S9 derivative with fluorescently labeled lp21 copies | (12) |

## References

1. Mead PS. 2015. Epidemiology of Lyme disease. Infect Dis Clin North Am 29:187– 210.

2. 2021. Lyme disease and relapsing fever spirochetes: genomics, molecular biology, host interactions and disease pathogenesis. Caister Academic Press, Norfolk.

3. Kugeler KJ, Earley A, Mead PS, Hinckley AF. 2024. Surveillance for Lyme Disease After Implementation of a Revised Case Definition - United States, 2022. MMWR Morb Mortal Wkly Rep 73:118–123.

4. Pustijanac E, Buršić M, Millotti G, Paliaga P, Iveša N, Cvek M. 2024. Tick-Borne Bacterial Diseases in Europe: Threats to public health. Eur J Clin Microbiol Infect Dis 10.1007/s10096-024-04836-5.

5. Bosler EM, Coleman JL, Benach JL, Massey DA, Hanrahan JP, Burgdorfer W, Barbour AG. 1983. Natural Distribution of the Ixodes dammini spirochete. Science 220:321–322.

6. Anderson JF, Duray PH, Magnarelli LA. 1987. Prevalence of Borrelia burgdorferi in white-footed mice and Ixodes dammini at Fort McCoy, Wis. J Clin Microbiol 25:1495–1497.

7. Anderson JF. 1989. Epizootiology of Borrelia in Ixodes tick vectors and reservoir hosts. Rev Infect Dis 11 Suppl 6:S1451–9.

8. Helble JD, McCarthy JE, Hu LT. 2021. Interactions between Borrelia burgdorferi and its hosts across the enzootic cycle. Parasite Immunol 43:e12816.

9. Oppler ZJ, O’Keeffe KR, McCoy KD, Brisson D. 2021. Evolutionary Genetics of Borrelia. Curr Issues Mol Biol 42:97–112.

10. Samuels DS, Lybecker MC, Yang XF, Ouyang Z, Bourret TJ, Boyle WK, Stevenson B, Drecktrah D, Caimano MJ. 2021. Gene Regulation and TranscriptomicsLyme Disease and Relapsing Fever Spirochetes: Genomics, Molecular Biology, Host Interactions and Disease Pathogenesis. Caister Academic Press.

11. Radolf JD, Caimano MJ, Stevenson B, Hu LT. 2012. Of ticks, mice and men: understanding the dual-host lifestyle of Lyme disease spirochaetes. Nat Rev Microbiol 10:87–99.

12. Steere AC, Strle F, Wormser GP, Hu LT, Branda JA, Hovius JWR, Li X, Mead PS. 2016. Lyme borreliosis. Nat Rev Dis Primers 2:16090.

13. Kung F, Anguita J, Pal U. 2013. Borrelia burgdorferi and tick proteins supporting pathogen persistence in the vector. Future Microbiol 8:41–56.

14. Caimano MJ, Drecktrah D, Kung F, Samuels DS. 2016. Interaction of the Lyme disease spirochete with its tick vector. Cell Microbiol 18:919–927.

15. Pal U, Kitsou C, Drecktrah D, Yaş ÖB, Fikrig E. 2021. Interactions between ticks and Lyme disease spirochetesLyme Disease and Relapsing Fever Spirochetes: Genomics, Molecular Biology, Host Interactions and Disease Pathogenesis. Caister Academic Press.

16. Piesman J, Oliver JR, Sinsky RJ. 1990. Growth kinetics of the Lyme disease spirochete (Borrelia burgdorferi) in vector ticks (Ixodes dammini). Am J Trop Med Hyg 42:352–357.

17. Piesman J, Schneider BS, Zeidner NS. 2001. Use of quantitative PCR to measure density of Borrelia burgdorferi in the midgut and salivary glands of feeding tick vectors. J Clin Microbiol 39:4145–4148.

18. Dunham-Ems SM, Caimano MJ, Eggers CH, Radolf JD. 2012. Borrelia burgdorferi requires the alternative sigma factor RpoS for dissemination within the vector during tick-to-mammal transmission. PLoS Pathog 8:e1002532.

19. Fazzino L, Tilly K, Dulebohn DP, Rosa PA. 2015. Long-term survival of Borrelia burgdorferi lacking the hibernation promotion factor homolog in the unfed tick vector. Infect Immun 83:4800–4810.

20. Stevenson B, Krusenstjerna AC, Castro-Padovani TN, Savage CR, Jutras BL, Saylor TC. 2022. The Consistent Tick-Vertebrate Infectious Cycle of the Lyme Disease Spirochete Enables Borrelia burgdorferi To Control Protein Expression by Monitoring Its Physiological Status. J Bacteriol 204:e0060621.

21. Tokarz R, Anderton JM, Katona LI, Benach JL. 2004. Combined effects of blood and temperature shift on Borrelia burgdorferi gene expression as determined by whole genome DNA array. Infect Immun 72:5419–5432.

22. Pappas CJ, Iyer R, Petzke MM, Caimano MJ, Radolf JD, Schwartz I. 2011. Borrelia burgdorferi requires glycerol for maximum fitness during the tick phase of the enzootic cycle. PLoS Pathog 7:e1002102.

23. Sonenshine DE, Michael Roe R. 2014. Biology of Ticks (Second Edition). Oxford University Press. Retrieved 18 June 2024.

24. Samanta K, Azevedo JF, Nair N, Kundu S, Gomes-Solecki M. 2022. Infected Ixodes scapularis Nymphs Maintained in Prolonged Questing under Optimal Environmental Conditions for One Year Can Transmit Borrelia burgdorferi (Borreliella genus novum) to Uninfected Hosts. Microbiol Spectr 10:e0137722.

25. Rosa PA, Tilly K, Stewart PE. 2005. The burgeoning molecular genetics of the Lyme disease spirochaete. Nat Rev Microbiol 3:129–143.

26. De Silva AM, Fikrig E. 1995. Growth and migration of Borrelia burgdorferi in Ixodes ticks during blood feeding. Am J Trop Med Hyg 53:397–404.

27. Piesman J, Schneider BS. 2002. Dynamic changes in Lyme disease spirochetes during transmission by nymphal ticks. Exp Appl Acarol 28:141–145.

28. Finkel SE. 2006. Long-term survival during stationary phase: evolution and the GASP phenotype. Nat Rev Microbiol 4:113–120.

29. Dworkin J, Harwood CS. 2022. Metabolic Reprogramming and Longevity in Quiescence. Annu Rev Microbiol 76:91–111.

30. Morita RY. 1988. Bioavailability of energy and its relationship to growth and starvation survival in nature. Can J Microbiol 34:436–441.

31. Hoehler TM, Jørgensen BB. 2013. Microbial life under extreme energy limitation. Nat Rev Microbiol 11:83–94.

32. Steinhaus EA, Birkeland JM. 1939. Studies on the Life and Death of Bacteria: I. The Senescent Phase in Aging Cultures and the Probable Mechanisms Involved. J Bacteriol 38:249–261.

33. Amy PS, Morita RY. 1983. Starvation-survival patterns of sixteen freshly isolated open-ocean bacteria. Appl Environ Microbiol 45:1109–1115.

34. Smeulders MJ, Keer J, Speight RA, Williams HD. 1999. Adaptation of Mycobacterium smegmatis to stationary phase. J Bacteriol 181:270–283.

35. Bergkessel M, Delavaine L. 2021. Diversity in Starvation Survival Strategies and Outcomes among Heterotrophic Proteobacteria. Microb Physiol 31:146–162.

36. Ratib NR, Seidl F, Ehrenreich IM, Finkel SE. 2021. Evolution in Long-Term Stationary-Phase Batch Culture: Emergence of Divergent Escherichia coli Lineages over 1,200 Days. MBio 12.

37. Nyström T. 2004. Stationary-phase physiology. Annu Rev Microbiol 58:161–181.

38. Fraser CM, Casjens S, Huang WM, Sutton GG, Clayton R, Lathigra R, White O, Ketchum KA, Dodson R, Hickey EK, Gwinn M, Dougherty B, Tomb JF, Fleischmann RD, Richardson D, Peterson J, Kerlavage AR, Quackenbush J, Salzberg S, Hanson M, van Vugt R, Palmer N, Adams MD, Gocayne J, Weidman J, Utterback T, Watthey L, McDonald L, Artiach P, Bowman C, Garland S, Fuji C, Cotton MD, Horst K, Roberts K, Hatch B, Smith HO, Venter JC. 1997. Genomic sequence of a Lyme disease spirochaete, Borrelia burgdorferi. Nature 390:580–586.

39. Kelly R. 1971. Cultivation of *Borrelia hermsi*. Science 173:443–444.

40. Barbour AG. 1984. Isolation and cultivation of Lyme disease spirochetes. Yale J Biol Med 57:521–525.

41. Elias AF, Bono JL, Carroll JA, Stewart P, Tilly K, Rosa P. 2000. Altered stationary-phase response in a Borrelia burgdorferi rpoS mutant. J Bacteriol 182:2909–2918.

42. Alban PS, Johnson PW, Nelson DR. 2000. Serum-starvation-induced changes in protein synthesis and morphology of Borrelia burgdorferi. Microbiology 146 (Pt 1):119–127.

43. Bugrysheva JV, Pappas CJ, Terekhova DA, Iyer R, Godfrey HP, Schwartz I, Cabello FC. 2015. Characterization of the RelBbu Regulon in Borrelia burgdorferi Reveals Modulation of Glycerol Metabolism by (p)ppGpp. PLoS One 10:e0118063.

44. Drecktrah D, Lybecker M, Popitsch N, Rescheneder P, Hall LS, Samuels DS. 2015. The Borrelia burgdorferi RelA/SpoT Homolog and Stringent Response Regulate Survival in the Tick Vector and Global Gene Expression during Starvation. PLoS Pathog 11:e1005160.

45. Meriläinen L, Herranen A, Schwarzbach A, Gilbert L. 2015. Morphological and biochemical features of Borrelia burgdorferi pleomorphic forms. Microbiology 161:516–527.

46. Drecktrah D, Hall LS, Rescheneder P, Lybecker M, Samuels DS. 2018. The Stringent Response-Regulated sRNA Transcriptome of Borrelia burgdorferi. Front Cell Infect Microbiol 8:231.

47. Boyle WK, Groshong AM, Drecktrah D, Boylan JA, Gherardini FC, Blevins JS, Samuels DS, Bourret TJ. 2019. DksA Controls the Response of the Lyme Disease Spirochete Borrelia burgdorferi to Starvation. J Bacteriol 201.

48. Feng J, Shi W, Zhang S, Zhang Y. 2015. Identification of new compounds with high activity against stationary phase Borrelia burgdorferi from the NCI compound collection. Emerg Microbes Infect 4:e31.

49. Sharma B, Brown AV, Matluck NE, Hu LT, Lewis K. 2015. Borrelia burgdorferi, the Causative Agent of Lyme Disease, Forms Drug-Tolerant Persister Cells. Antimicrob Agents Chemother 59:4616–4624.

50. Feng J, Shi W, Zhang S, Sullivan D, Auwaerter PG, Zhang Y. 2016. A Drug Combination Screen Identifies Drugs Active against Amoxicillin-Induced Round Bodies of In Vitro Borrelia burgdorferi Persisters from an FDA Drug Library. Front Microbiol 7:743.

51. Cabello FC, Embers ME, Newman SA, Godfrey HP. 2022. Borreliella burgdorferi Antimicrobial-Tolerant Persistence in Lyme Disease and Posttreatment Lyme Disease Syndromes. MBio 13:e0344021.

52. Takacs CN, Wachter J, Xiang Y, Ren Z, Karaboja X, Scott M, Stoner MR, Irnov I, Jannetty N, Rosa PA, Wang X, Jacobs-Wagner C. 2022. Polyploidy, regular patterning of genome copies, and unusual control of DNA partitioning in the Lyme disease spirochete. Nat Commun 13:7173.

53. Iyer R, Mukherjee P, Wang K, Simons J, Wormser GP, Schwartz I. 2013. Detection of Borrelia burgdorferi nucleic acids after antibiotic treatment does not confirm viability. J Clin Microbiol 51:857–862.

54. Hinnebusch J, Barbour AG. 1992. Linear- and circular-plasmid copy numbers in Borrelia burgdorferi. J Bacteriol 174:5251–5257.

55. Sădziene A, Rosa PA, Thompson PA, Hogan DM, Barbour AG. 1992. Antibody- resistant mutants of Borrelia burgdorferi: in vitro selection and characterization. J Exp Med 176:799–809.

56. Fulton JD, Smith PJC. 1960. Carbohydrate metabolism in Spirochaeta recurrentis. 1. The metabolism of spirochaetes in vivo and in vitro. Biochem J 76:491–499.

57. Smith PJ. 1960. Carbohydrate metabolism in Spirochaeta recurrentis. 2. Enzymes associated with disintegrated cells and extracts of spirochaetes. Biochem J 76:500–508.

58. Barbour AG, Hayes SF. 1986. Biology of Borrelia species. Microbiol Rev 50:381– 400.

59. Rosa PA. 1997. Microbiology of Borrelia burgdorferi. Semin Neurol 17:5–10.

60. Caimano MJ, Eggers CH, Hazlett KRO, Radolf JD. 2004. RpoS is not central to the general stress response in Borrelia burgdorferi but does control expression of one or more essential virulence determinants. Infect Immun 72:6433–6445.

61. Yang X, Goldberg MS, Popova TG, Schoeler GB, Wikel SK, Hagman KE, Norgard MV. 2000. Interdependence of environmental factors influencing reciprocal patterns of gene expression in virulent Borrelia burgdorferi. Mol Microbiol 37:1470–1479.

62. Brorson O, Brorson SH. 1997. Transformation of cystic forms of Borrelia burgdorferi to normal, mobile spirochetes. Infection 25:240–246.

63. Drecktrah D, Hall LS, Crouse B, Schwarz B, Richards C, Bohrnsen E, Wulf M, Long B, Bailey J, Gherardini F, Bosio CM, Lybecker MC, Samuels DS. 2022. The glycerol-3-phosphate dehydrogenases GpsA and GlpD constitute the oxidoreductive metabolic linchpin for Lyme disease spirochete host infectivity and persistence in the tick. PLoS Pathog 18:e1010385.

64. Jutras BL, Scott M, Parry B, Biboy J, Gray J, Vollmer W, Jacobs-Wagner C. 2016. Lyme disease and relapsing fever *Borrelia* elongate through zones of peptidoglycan synthesis that mark division sites of daughter cells. Proc Natl Acad Sci U S A 113:9162–9170.

65. Hyde FW, Johnson RC. 1986. Genetic analysis of Borrelia. Zentralbl Bakteriol Mikrobiol Hyg A 263:119–122.

66. Barbour AG. 1988. Plasmid analysis of Borrelia burgdorferi, the Lyme disease agent. J Clin Microbiol 26:475–478.

67. Schwan TG, Burgdorfer W, Garon CF. 1988. Changes in infectivity and plasmid profile of the Lyme disease spirochete, Borrelia burgdorferi, as a result of in vitro cultivation. Infect Immun 56:1831–1836.

68. Norris SJ, Carter CJ, Howell JK, Barbour AG. 1992. Low-passage-associated proteins of Borrelia burgdorferi B31: characterization and molecular cloning of OspD, a surface-exposed, plasmid-encoded lipoprotein. Infect Immun 60:4662– 4672.

69. Norris SJ, Howell JK, Garza SA, Ferdows MS, Barbour AG. 1995. High- and low-infectivity phenotypes of clonal populations of in vitro-cultured Borrelia burgdorferi. Infect Immun 63:2206–2212.

70. Purser JE, Norris SJ. 2000. Correlation between plasmid content and infectivity in Borrelia burgdorferi. Proc Natl Acad Sci U S A 97:13865–13870.

71. Bunikis I, Kutschan-Bunikis S, Bonde M, Bergström S. 2011. Multiplex PCR as a tool for validating plasmid content of Borrelia burgdorferi. J Microbiol Methods 86:243–247.

72. Indest KJ, Ramamoorthy R, Solé M, Gilmore RD, Johnson BJ, Philipp MT. 1997. Cell-density-dependent expression of Borrelia burgdorferi lipoproteins in vitro. Infect Immun 65:1165–1171.

73. Kurokawa C, Lynn GE, Pedra JHF, Pal U, Narasimhan S, Fikrig E. 2020. Interactions between Borrelia burgdorferi and ticks. Nat Rev Microbiol 18:587–600.

74. Kell DB, Kaprelyants AS, Weichart DH, Harwood CR, Barer MR. 1998. Viability and activity in readily culturable bacteria: a review and discussion of the practical issues. Antonie Van Leeuwenhoek 73:169–187.

75. Bogosian G, Bourneuf EV. 2001. A matter of bacterial life and death. EMBO Rep 2:770–774.

76. Byram R, Stewart PE, Rosa P. 2004. The essential nature of the ubiquitous 26-kilobase circular replicon of Borrelia burgdorferi. J Bacteriol 186:3561–3569.

77. Finkel SE, Kolter R. 1999. Evolution of microbial diversity during prolonged starvation. Proc Natl Acad Sci U S A 96:4023–4027.

78. Gray DA, Dugar G, Gamba P, Strahl H, Jonker MJ, Hamoen LW. 2019. Extreme slow growth as alternative strategy to survive deep starvation in bacteria. Nat Commun 10:890.

79. Smith TC 2nd, Helm SM, Chen Y, Lin Y-H, Rajasekhar Karna SL, Seshu J. 2018. Borrelia host adaptation protein (BadP) is required for the colonization of a mammalian host by the agent of Lyme disease. Infect Immun 86.

80. Barker RJ, Lehner Y. 1976. Sugars in hemolymph of ticks. J Med Entomol 13:379– 380.

81. Tilly K, Grimm D, Bueschel DM, Krum JG, Rosa P. 2004. Infectious cycle analysis of a Borrelia burgdorferi mutant defective in transport of chitobiose, a tick cuticle component. Vector Borne Zoonotic Dis 4:159–168.

82. von Lackum K, Stevenson B. 2005. Carbohydrate utilization by the Lyme borreliosis spirochete, Borrelia burgdorferi. FEMS Microbiol Lett 243:173–179.

83. Wickramasekara S, Bunikis J, Wysocki V, Barbour AG. 2008. Identification of residual blood proteins in ticks by mass spectrometry proteomics. Emerg Infect Dis 14:1273–1275.

84. He M, Ouyang Z, Troxell B, Xu H, Moh A, Piesman J, Norgard MV, Gomelsky M, Yang XF. 2011. Cyclic di-GMP is essential for the survival of the lyme disease spirochete in ticks. PLoS Pathog 7:e1002133.

85. Hoon-Hanks LL, Morton EA, Lybecker MC, Battisti JM, Samuels DS, Drecktrah D. 2012. Borrelia burgdorferi malQ mutants utilize disaccharides and traverse the enzootic cycle. FEMS Immunol Med Microbiol 66:157–165.

86. Laskay UA, Breci L, Vilcins I-ME, Dietrich G, Barbour AG, Piesman J, Wysocki VH. 2013. Survival of host blood proteins in Ixodes scapularis (Acari: Ixodidae) ticks: a time course study. J Med Entomol 50:1282–1290.

87. Caimano MJ, Dunham-Ems S, Allard AM, Cassera MB, Kenedy M, Radolf JD. 2015. Cyclic di-GMP modulates gene expression in Lyme disease spirochetes at the tick-mammal interface to promote spirochete survival during the blood meal and tick-to-mammal transmission. Infect Immun 83:3043–3060.

88. Corona A, Schwartz I. 2015. Borrelia burgdorferi: Carbon Metabolism and the Tick-Mammal Enzootic Cycle. Microbiol Spectr 3.

89. Sonenshine DE, Šimo L. 2021. Biology and Molecular Biology of *Ixodes scapularis*, p. 339–366. In Justin D. Radolf, (ed)., D. Scott Samuels, (ed). (eds.), Lyme Disease and Relapsing Fever Spirochetes: Genomics, Molecular Biology, Host Interactions and Disease Pathogenesis.

90. Schink SJ, Biselli E, Ammar C, Gerland U. 2019. Death Rate of E. coli during Starvation Is Set by Maintenance Cost and Biomass Recycling. Cell Syst 9:64–73.e3.

91. Sidak-Loftis LC, Rosche KL, Pence N, Ujczo JK, Hurtado J, Fisk EA, Goodman AG, Noh SM, Peters JW, Shaw DK. 2022. The Unfolded-Protein Response Triggers the Arthropod Immune Deficiency Pathway. MBio 13:e0070322.

92. Stiefel P, Schmidt-Emrich S, Maniura-Weber K, Ren Q. 2015. Critical aspects of using bacterial cell viability assays with the fluorophores SYTO9 and propidium iodide. BMC Microbiol 15:36.

93. Rosenberg M, Azevedo NF, Ivask A. 2019. Propidium iodide staining underestimates viability of adherent bacterial cells. Sci Rep 9:6483.

94. Norris SJ, Howell JK, Odeh EA, Lin T, Gao L, Edmondson DG. 2011. High-throughput plasmid content analysis of Borrelia burgdorferi B31 by using Luminex multiplex technology. Appl Environ Microbiol 77:1483–1492.

95. Samuels DS. 1995. Electrotransformation of the spirochete Borrelia burgdorferi. Methods Mol Biol 47:253–259.

96. Elias AF, Stewart PE, Grimm D, Caimano MJ, Eggers CH, Tilly K, Bono JL, Akins DR, Radolf JD, Schwan TG, Rosa P. 2002. Clonal polymorphism of Borrelia burgdorferi strain B31 MI: implications for mutagenesis in an infectious strain background. Infect Immun 70:2139–2150.

97. Stewart PE, Rosa PA. 2008. Transposon Mutagenesis of the Lyme Disease AgentBorrelia burgdorferi, p. 85–95. In DeLeo, FR, Otto, M (eds.), Bacterial Pathogenesis: Methods and Protocols. Humana Press, Totowa, NJ.

98. Samuels DS, Drecktrah D, Hall LS. 2018. Genetic Transformation and Complementation. Methods Mol Biol 1690:183–200.

99. Seshu J, Moy BE, Ingle TM. 2021. Transformation of Borrelia burgdorferi. Curr Protoc 1:e61.

100. Lanz MC, Zatulovskiy E, Swaffer MP, Zhang L, Ilerten I, Zhang S, You DS, Marinov G, McAlpine P, Elias JE, Skotheim JM. 2022. Increasing cell size remodels the proteome and promotes senescence. Mol Cell 82:3255–3269.e8.

101. Lanz MC, Zhang S, Swaffer MP, Ziv I, Götz LH, McCarty F, Jarosz DF, Elias JE, Skotheim JM. 2023. Genome dilution by cell growth drives starvation-like proteome remodeling in mammalian and yeast cells. bioRxiv.

102. Mäkelä J, Papagiannakis A, Lin W-H, Lanz MC, Glenn S, Swaffer M, Marinov GK, Skotheim JM, Jacobs-Wagner C. 2024. Genome concentration limits cell growth and modulates proteome composition in Escherichia coli. eLife.

103. Zückert WR. 2007. Laboratory maintenance of Borrelia burgdorferi. Curr Protoc Microbiol Chapter 12:Unit 12C.1.

104. Jutras BL, Chenail AM, Stevenson B. 2013. Changes in bacterial growth rate govern expression of the Borrelia burgdorferi OspC and Erp infection-associated surface proteins. J Bacteriol 195:757–764.

105. Takacs CN, Scott M, Chang Y, Kloos ZA, Irnov I, Rosa PA, Liu J, Jacobs-Wagner C. 2021. A CRISPR interference platform for selective downregulation of gene expression in Borrelia burgdorferi. Appl Environ Microbiol 87.

106. Lindenbach BD, Evans MJ, Syder AJ, Wölk B, Tellinghuisen TL, Liu CC, Maruyama T, Hynes RO, Burton DR, McKeating JA, Rice CM. 2005. Complete replication of hepatitis C virus in cell culture. Science 309:623–626.

107. Reed LJ, Muench H. 1938. A SIMPLE METHOD OF ESTIMATING FIFTY PER CENT ENDPOINTS. Am J Epidemiol 27:493–497.

108. Glaser P, Sharpe ME, Raether B, Perego M, Ohlsen K, Errington J. 1997. Dynamic, mitotic-like behavior of a bacterial protein required for accurate chromosome partitioning. Genes Dev 11:1160–1168.

109. Paintdakhi A, Parry B, Campos M, Irnov I, Elf J, Surovtsev I, Jacobs-Wagner C. 2016. Oufti: an integrated software package for high-accuracy, high-throughput quantitative microscopy analysis. Mol Microbiol 99:767–777.

110. Otsu N. 1979. A Threshold Selection Method from Gray-Level Histograms. IEEE Trans Syst Man Cybern 9:62–66.

111. Rego ROM, Bestor A, Rosa PA. 2011. Defining the plasmid-borne restriction-modification systems of the Lyme disease spirochete Borrelia burgdorferi. J Bacteriol 193:1161–1171.

112. McKinney W. 2010. Data Structures for Statistical Computing in Python, p. 56–61. In Proceedings of the Python in Science Conference. SciPy.

113. The pandas development team. 2024. pandas-dev/pandas: Pandas. Zenodo. https://zenodo.org/records/10957263. Retrieved 4 September 2024.

114. Schindelin J, Arganda-Carreras I, Frise E, Kaynig V, Longair M, Pietzsch T, Preibisch S, Rueden C, Saalfeld S, Schmid B, Tinevez J-Y, White DJ, Hartenstein V, Eliceiri K, Tomancak P, Cardona A. 2012. Fiji: an open-source platform for biological-image analysis. Nat Methods 9:676–682.

## References

1. Burgdorfer W, Barbour AG, Hayes SF, Benach JL, Grunwaldt E, Davis JP. 1982. Lyme disease-a tick-borne spirochetosis? Science 216:1317–1319.

2. Fraser CM, Casjens S, Huang WM, Sutton GG, Clayton R, Lathigra R, White O, Ketchum KA, Dodson R, Hickey EK, Gwinn M, Dougherty B, Tomb JF, Fleischmann RD, Richardson D, Peterson J, Kerlavage AR, Quackenbush J, Salzberg S, Hanson M, van Vugt R, Palmer N, Adams MD, Gocayne J, Weidman J, Utterback T, Watthey L, McDonald L, Artiach P, Bowman C, Garland S, Fuji C, Cotton MD, Horst K, Roberts K, Hatch B, Smith HO, Venter JC. 1997. Genomic sequence of a Lyme disease spirochaete, Borrelia burgdorferi. Nature 390:580–586.

3. Casjens S, Palmer N, van Vugt R, Huang WM, Stevenson B, Rosa P, Lathigra R, Sutton G, Peterson J, Dodson RJ, Haft D, Hickey E, Gwinn M, White O, Fraser CM. 2000. A bacterial genome in flux: the twelve linear and nine circular extrachromosomal DNAs in an infectious isolate of the Lyme disease spirochete Borrelia burgdorferi. Mol Microbiol 35:490–516.

4. Barthold SW, Moody KD, Terwilliger GA, Duray PH, Jacoby RO, Steere AC. 1988. Experimental Lyme arthritis in rats infected with Borrelia burgdorferi. J Infect Dis 157:842–846.

5. Barthold SW, Moody KD, Terwilliger GA, Jacoby RO, Steere AC. 1988. An animal model for Lyme arthritis. Ann N Y Acad Sci 539:264–273.

6. Steere AC, Grodzicki RL, Kornblatt AN, Craft JE, Barbour AG, Burgdorfer W, Schmid GP, Johnson E, Malawista SE. 1983. The spirochetal etiology of Lyme disease. N Engl J Med 308:733–740.

7. Steere AC, Grodzicki RL, Craft JE, Shrestha M, Kornblatt AN, Malawista SE. 1984. Recovery of Lyme disease spirochetes from patients. Yale J Biol Med 57:557–560.

8. Schutzer SE, Fraser-Liggett CM, Casjens SR, Qiu W-G, Dunn JJ, Mongodin EF, Luft BJ. 2011. Whole-genome sequences of thirteen isolates of Borrelia burgdorferi. J Bacteriol 193:1018–1020.

9. Casjens SR, Mongodin EF, Qiu W-G, Luft BJ, Schutzer SE, Gilcrease EB, Huang WM, Vujadinovic M, Aron JK, Vargas LC, Freeman S, Radune D, Weidman JF, Dimitrov GI, Khouri HM, Sosa JE, Halpin RA, Dunn JJ, Fraser CM. 2012. Genome stability of Lyme disease spirochetes: comparative genomics of Borrelia burgdorferi plasmids. PLoS One 7:e33280.

10. Dunham-Ems SM, Caimano MJ, Pal U, Wolgemuth CW, Eggers CH, Balic A, Radolf JD. 2009. Live imaging reveals a biphasic mode of dissemination of Borrelia burgdorferi within ticks. J Clin Invest 119:3652–3665.

11. Rego ROM, Bestor A, Rosa PA. 2011. Defining the plasmid-borne restriction-modification systems of the Lyme disease spirochete Borrelia burgdorferi. J Bacteriol 193:1161–1171.

12. Takacs CN, Wachter J, Xiang Y, Ren Z, Karaboja X, Scott M, Stoner MR, Irnov I, Jannetty N, Rosa PA, Wang X, Jacobs-Wagner C. 2022. Polyploidy, regular patterning of genome copies, and unusual control of DNA partitioning in the Lyme disease spirochete. Nat Commun 13:7173.

